# Sequential Sampling as a Computational Principle of Episodic Memory

**DOI:** 10.64898/2026.10.08.757704

**Authors:** Casper Kerrén, Shrey Dixit, Evie Coxon, Andrey R Nikolaev, Christian F Doeller

## Abstract

Visual perception unfolds through sequential eye movements, yet computational models of episodic memory typically assume that event representations already exist. Here, we propose that sequential gaze sampling does not simply supply memory with visual information, but helps construct the representations that are subsequently stored and retrieved. We introduce a computational model in which successive fixation-dependent samples are integrated into an evolving internal representation before one-shot storage in associative memory. Within individual elements, successive samples progressively individuate the representation; across elements, transitions promote relational integration. Crucially, both effects arise from the same history-dependent updating process where each new sample is integrated into a state already shaped by what came before. As a result, sampling history becomes embedded in the memory representation itself. Changing the order of otherwise identical samples alters both the resulting representation and element-specific retrieval, while reinstating the original sampling trajectory facilitates access to the stored memory. Model-neural analysis further show that changes in the model’s evolving representations correspond to neural representational dynamics in lateral temporal cortex. Together, these findings support sequential sampling as a candidate computational process for constructing representations available to episodic memory, linking ongoing perceptual sampling to subsequent memory formation and retrieval.

## Introduction

Episodic memory allows us to remember specific events from our personal past, including what happened, where, and when ^1,2^. Yet such events are not encountered as complete snapshots. During natural viewing, the eyes sample only a small part of the visual environment at any moment, so an event must be constructed across a sequence of partial observations, accumulating information about individual elements and linking them across space and time. But how does the brain transform sequential gaze samples into an episodic memory?

Eye movements play multiple roles in episodic memory. During encoding, repeated sampling within an element supports later recognition ^3,4^, whereas transitions between elements may help establish the relations that define a coherent event representation ^5–8^. At retrieval, restricting eye movements impairs recognition ^9^, gaze patterns can reveal memory for associations without explicit reports ^10^, and retrieval can be facilitated by reinstating gaze sequences established during encoding ^11–16^. These functions are linked to the hippocampal memory system and its interactions with oculomotor and cortical perceptual systems ^17–24^. Yet perceptual information is progressively transformed across the cortical hierarchy ^25–27^, with the hippocampus receiving highly transformed cortical input rather than a direct copy of the visual environment ^28–30^.

Visual information acquired during successive fixations can be integrated across saccades within transsaccadic visual or short-term memory representations ^31–36^, potentially providing an intermediate stage between perceptual input and episodic storage. An important question is whether the temporal organisation of this integration shapes the representations subsequently available to episodic memory. We propose that it does: sustained sampling within an element progressively individuates that element, whereas transitions between elements integrate information across them. The organisation of gaze may therefore determine the relative contributions of element individuation and relational integration to the resulting representation

This role of sequential sampling and particularly how sampling is organised, is largely missing from current computational theories of episodic memory. Many existing memory models begin with an already formed representation and ask how it is stored, associated, consolidated, or retrieved ^37–43^. Temporal-context models explain how temporal history shapes memory by allowing encountered or retrieved items to update an evolving context state that supports the temporal organisation of retrieval ^44,45^. These principles extend to visual exploration, where reinstatement of fixation sequences exhibits temporal contiguity and is associated with successful memory ^46^. However, these accounts do not explicitly specify how successive fixation-dependent visual samples construct the representation subsequently available for episodic storage.

Meanwhile, models of visual sampling primarily address perception, attention or oculomotor behaviour ^47–51^. Recent work shows that recurrent prediction of successive fixations can construct integrated scene representations aligned with human visual cortex ^52^, but does not address how a particular sampling trajectory is stored as a novel episodic memory after a single encounter. Related frameworks connect fixation-level information extraction to event-model construction across multiple fixations in working memory and episodic long-term memory ^53^, but do not specify how sampling history transforms the element-specific representation available for associative storage and retrieval. Models explicitly linking sequential eye movements to memory have likewise focused primarily on visual recognition ^54,55^. Thus, an important computational question remains: how does a sequence of partial perceptual samples become the representation subsequently available to episodic memory?

In our model, successive fixations are integrated over time, such that each new sample contributes to an evolving representation. This account makes four predictions. First, within an element, additional observations should progressively reduce uncertainty and differentiate that element from similar ones, with semantic information becoming available before the information needed to distinguish that particular element ^34,56–59^. This increasing differentiation may provide the representational input on which hippocampal pattern separation subsequently operates ^60^. Second, when sampling moves between elements, their representations are integrated within the same unfolding sequence ^3,5–7,22^. Third, as successive samples are integrated, the evolving representation should become organised along fewer dominant dimensions while retaining information about where, when, and in what order observations occurred ^7,61–63^, and changing temporal order should therefore alter element-specific representations. Finally, because sampling history remains embedded in the representation, retrieval should benefit from reinstating the trajectory through which an event was encoded, with element-specific memory depending more strongly on this reinstatement than semantic information ^14,15,64^. Such reinstatement may provide a route to hippocampal pattern completion ^64–66^. Sequential sampling may therefore shape the representations that distinguish individual elements, the relations that connect them within an episode, and the cues through which those representations are subsequently retrieved.

To test these predictions, we developed a model in which gaze selects local visual representations that are sequentially integrated and stored in associative memory (Fig. 1). The model separates sequential integration from associative storage: successive samples construct an evolving representation whose final state is stored in associative memory. By manipulating the amount, location, temporal order, and between-element organisation of sampling, we test how sequential gaze shapes element individuation and relational binding, representational geometry, and retrieval from partial cues. Together, these analyses test whether the representation available to episodic memory is determined not only by what information is sampled, but by the trajectory through which it is acquired.

**Figure 1.**
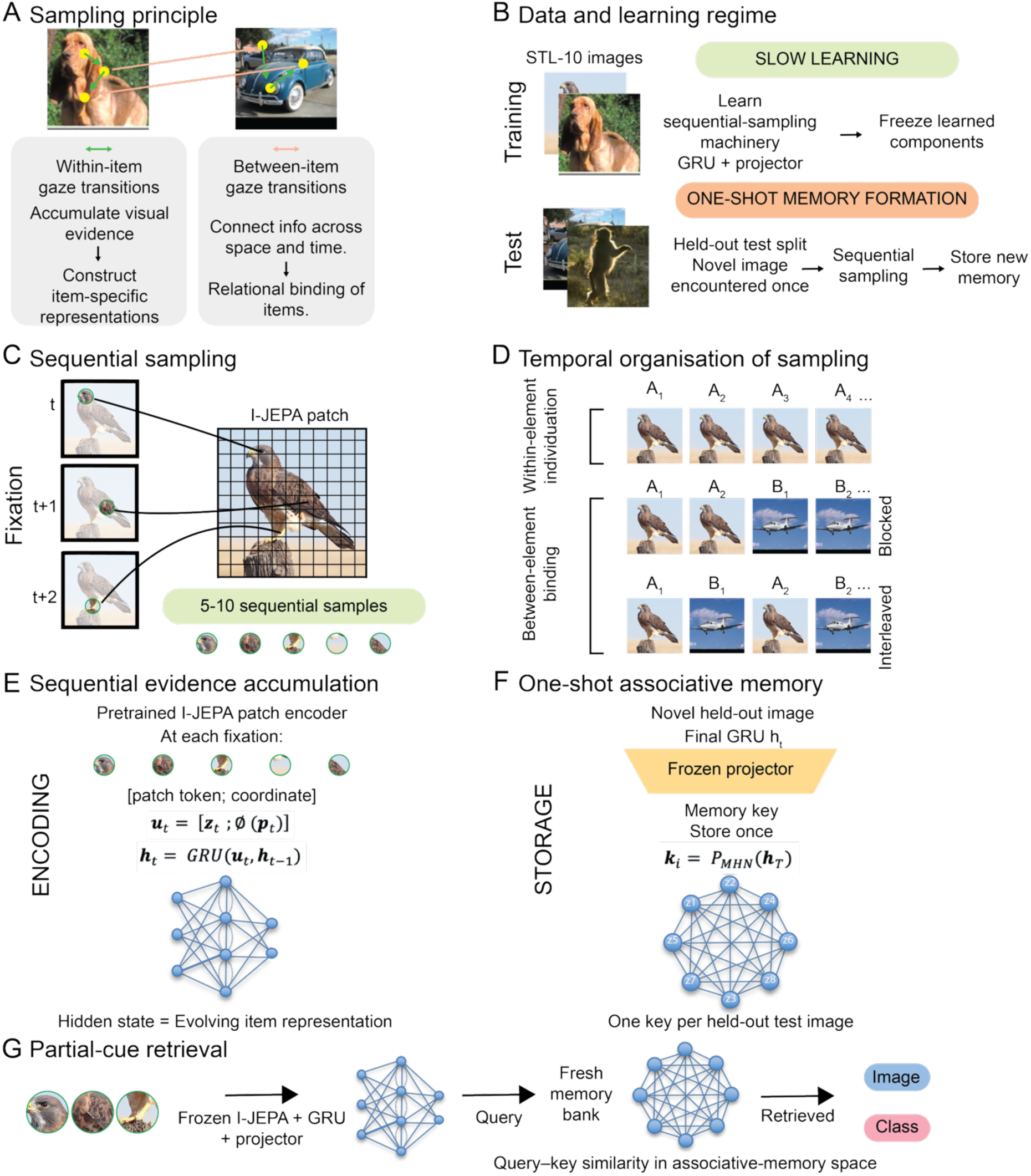
Computational framework for sequential sampling and episodic memory. **A** Sequential sampling is proposed to support both individuation within elements and relational binding across elements. B The recurrent and projection networks are learned using the STL-10 training split and subsequently frozen. Novel held-out test images are then encoded once and stored as new memories without further parameter updates, separating slow learning of the sampling mechanism from one-shot memory formation. C Each element is represented by a sequence of spatially localised l-JEPA patch representations selected by successive fixations. D The temporal organisation of otherwise identical samples is manipulated to test within-item individuation and between-item relational binding. E A GRU recurrently integrates visual and spatial information across successive samples to construct an evolving element representation. F The final recurrent state is mapped through the frozen projector and stored as a new key in a Modern Hopfield Network (MHN). G At retrieval, partial or perturbed sampling trajectories are processed through the same frozen pathway to generate queries to the newly formed memory bank, allowing image-specific and semantic retrieval to be assessed.

### Computational framework

An overview of the model can be found in Figure 1. We modelled individual natural images from the STL-10 dataset ^67^, treating each image as a distinct element belonging to one of ten semantic categories. Rather than presenting an entire image to the memory system at once, visual input was represented as a sequence of fixation-dependent local samples.

Each within-element fixation selected a spatially local patch representation from a pretrained I-JEPA visual model and was accompanied by its fixation coordinates. These inputs were presented sequentially to a gated recurrent unit (GRU), whose hidden state was updated after each sample. The GRU was trained on STL-10 training images using a next-fixation prediction objective: from preceding samples, it predicted the visual representation encountered at the subsequent fixation. This encouraged the recurrent state to integrate accumulated visual, spatial, and temporal information rather than representing each glimpse independently. For the primary encoding procedure, each image was represented by a centre-biased, variable-length trajectory of five to ten fixations; analyses requiring matched observations at every serial position used complete ten-fixation trajectories.

After GRU training, its parameters were frozen. A separate projection network was trained on the training split to map recurrent states into associative-memory space. During projector training, states generated from partial fixation sequences were aligned with the projected final state of the same sampled trajectory, while remaining discriminable from other training images. The projected final state provided the key representation used for associative storage, whereas projected partial states provided retrieval queries.

Unless otherwise specified, all reported analyses used a held-out one-shot memory procedure. The pretrained I-JEPA encoder, GRU, and projection network were applied without further updating to STL-10 test images. For each analysis, a new memory bank was formed by storing one projected final state for each held-out image. Partial, degraded, or temporally perturbed fixation sequences from those same test images were then processed through the frozen pathway and compared with this newly formed test-image memory bank. Exact-image performance therefore measured retrieval of newly stored elements after a single encoding experience, rather than recognition of images used to train the recurrent or projection networks.

For between-element fixations supporting relational memory, we used the same frozen components and held-out test images without additional model learning. In the heteroassociative analysis, newly formed A representations were stored as keys associated with newly formed B representations as values. In the temporal-history analysis, fixation samples from A and B were integrated continuously into one joint recurrent sequence, and its final projected state was stored as a joint A-B memory representation. Thus, identical samples could be presented with different temporal organisation while holding the learned visual and recurrent pathways fixed.

At retrieval, a partial fixation sequence was passed through the frozen GRU and projection pathway to generate a query to the Modern Hopfield Network (MHN). Queries and stored keys were L2-normalised, and their cosine similarities determined retrieval weights over the stored memories; the highest-scoring key defined top-1 retrieval. This architecture allowed us to measure how the evolving representation approached the final stored state and how the amount, order, and reinstatement of sampled information affected memory access.

Separate generative analyses explored downstream consequences of the resulting memory representations, including fixation-constrained reconstruction. These readout models were not required for associative storage or retrieval.

## Results

### Sequential sampling progressively constructs element-specific memory representations

Our central hypothesis is that representations available to episodic memory are not formed from complete perceptual inputs, but are constructed gradually as information is sampled over time. We therefore first asked whether each additional fixation makes the identity of an individual element less uncertain.

Each image was represented as a sequence of local visual samples integrated into an evolving recurrent state (Fig. 2A-B). After each fixation, the current state was projected into memory space and used to query the fixed memory bank, allowing retrieval evidence and uncertainty to be tracked across successive samples.

**Figure 2.**
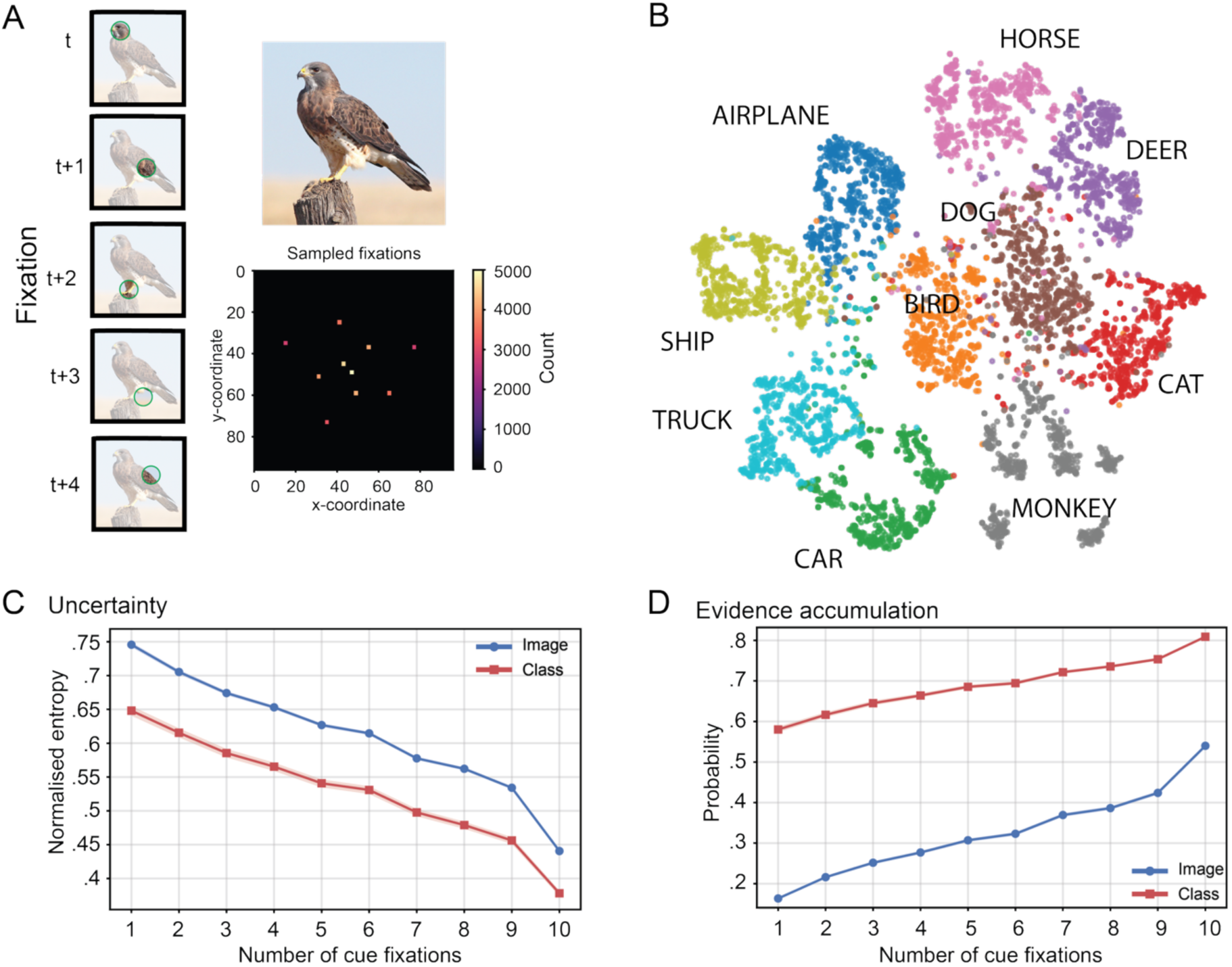
Sequential sampling progressively constructs element-specific memory representations. **A.** Visual input is encoded through a sequence of attended fixation locations rather than as a complete image. Each fixation is mapped to its corresponding l-JEPA patch representation, producing an ordered sequence of visual samples. Variable fixation trajectories (5-10 fixations) are used to model natural variability in gaze sampling, while the distribution of sampled fixation locations across the entire training set is shown below. B. The resulting episodic memory representations form a structured latent space in which semantic categories form distinct clusters despite memories being constructed from sequential samples. C. As additional fixations are accumulated, representational uncertainty progressively decreases Both image-level and class-level entropy decline across successive fixation prefixes, reflecting increasingly stable episodic representations. D. Evidence accumulates throughout sequential sampling The probability assigned to the correct semantic class rises rapidly and precedes the slower accumulation of evidence supporting the correct image identity, indicating that semantic structure emerges before precise episodic identity.

Successive fixations progressively reduced uncertainty about both the specific image and its semantic category (Fig. 2C). Normalised image-level entropy decreased from 0.746 (95% bootstrap CI [0.742, 0.749]) after the first fixation to 0.441 [0.439, 0.442] after ten fixations, while class-level entropy decreased from 0.648 [0.641, 0.655] to 0.378 [0.374, 0.382], both bootstrap p <.001. Thus, additional samples progressively narrowed the set of memories consistent with the observed evidence ^68^.

Consistent with this reduction in uncertainty, evidence for the correct stored memory accumulated across the sequence (Fig. 2D). Target-image probability increased from 0.163 [0.161, 0.166] after one fixation to 0.540 [0.536, 0.544] after ten fixations, while target-class probability increased from 0.580 [0.573, 0.587] to 0.809 [0.807, 0.812], both bootstrap p <.001. The target’s margin over its nearest same-class competitor also increased from 0.169 [0.165, 0.173] to 0.327 [0.322, 0.331], bootstrap p <.001. This accumulation was amplified after projection into associative-memory space relative to the raw recurrent state (Supplementary Fig. 1).

A complementary raw-state analysis assessed when the correct class or exact image became stably discriminable. Stable class discriminability emerged after 1.26 fixations [1.21, 1.32], and stable element discriminability after 1.58 fixations [1.52, 1.64], bootstrap p <.001. Thus, both semantic and image-specific evidence became available rapidly, although semantic evidence stabilised slightly earlier.

Together, these findings show how an individuated memory representation can be constructed from sequentially sampled perceptual evidence. Each fixation reduced uncertainty and strengthened element-specific evidence, with broad semantic structure becoming discriminable before precise element identity.

### Sequential sampling supports relational binding across elements

The preceding analyses showed how successive samples progressively construct an individuated representation of a single element. Episodic memory, however, also requires information encountered across an experience to become relationally bound, such that one element can subsequently provide access to another. Eye movements are particularly sensitive to such relational memory, with viewing behaviour expressing learned relationships among elements even in the absence of explicit reports of remembering ^10^. We therefore asked how both the amount and temporal organisation of sampling across elements determine subsequent relational accessibility.

We first tested whether accumulating fixation-derived information about one element improved access to its associate. Using arbitrary pairs of held-out STL-10 images, final A representations were stored as MHN keys and paired B representations as values (Fig. 3A). As more fixation evidence from A became available, retrieval of its paired B improved systematically (Fig. 3B), increasing from 39.8% [38.0%, 41.5%] with one fixation to 62.4% [60.8%, 64.0%] with five fixations, bootstrap p <.001; chance was 4.2%. This advantage was not explained simply by sampled regions being more informative about A itself: after matching sampled and unsampled regions for A diagnosticity, sampled regions still assigned greater probability to the correct B, 0.242 [0.232, 0.251] versus 0.210 [0.201, 0.219], bootstrap p <.001 (Fig. 3C).

**Figure 3.**
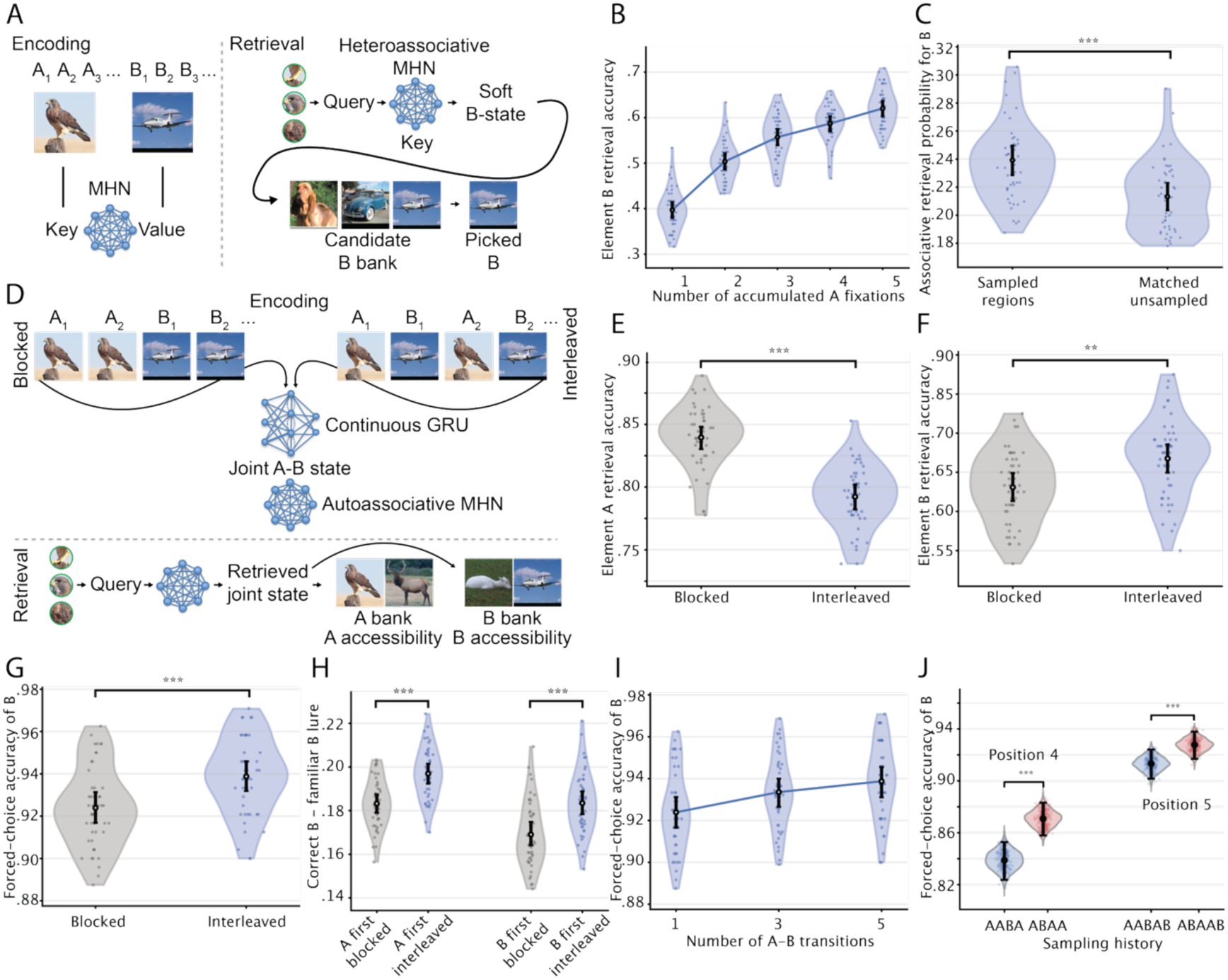
Sequential sampling supports relational binding across event elements. **A** Schematic of the heteroassociative A-to-B retrieval analysis. Final representations of A elements were stored as MHN keys and their paired B representations as values. Fixation-derived cues from A produced a soft B-valued state, which was compared with the candidate B representation bank to identify the retrieved associate. B Accumulating more fixation evidence from A progressively increased access to its assigned associate B. C We compared individual regions sampled during A-B encoding with unsampled regions matched for their diagnosticity of A. Sampled regions assigned greater probability to the correct associate than diagnosticity-matched unsampled regions. D Schematic of the temporal-organisation analysis. The same fixation samples from A and B were presented in blocked (AAABBB) or interleaved (ABABAB) sequencesand integrated continuously by the GRU without resetting between elements. The projected final state formed a joint A-B representation stored in an autoassociative MHN. At retrieval, an A-derived cue recovered the joint state, whose similarity to separate A and B representation banks quantified the accessibility of each element. E Retrieval accuracy for cue element A following blocked and interleaved encoding. F Retrieval accuracy for associated item B following blocked and interleaved encoding. G Forced-choice retrieval of the correct B against a familiarity-matched B lure was higher following interleaved than blocked encoding. H The similarity advantage of the correct B over its matched lure was greater following interleaved encoding, regardless of whetherthe sequence began with A or B, I Forced-choice retrieval of the correct B increased progressively with the number of A-B transitions during encoding. J Correct-B forced-choice accuracy was higher following ABAA than AABA sampling at position 4 and following ABAAB than AABAB at position 5, despite matched visual samples and transition counts within each position. Violins and points show bootstrap estimates, and black markers and error bars indicate observed accuracy and bootstrap 95% confidence intervals.

We next tested our central prediction that temporal transitions between elements would strengthen relational binding. Fixation samples from A and B were integrated within a single continuous sequence without resetting the GRU, and the final projected state was stored as a joint A-B representation (Fig. 3D). We compared blocked sequences, in which three different samples from each element occurred consecutively, with interleaved sequences that repeatedly alternated between A and B. If transitions between elements contribute to relational binding ^5–8^, interleaved sampling should strengthen access to B, potentially at the expense of maintaining a more distinct representation of A. Exact A retrieval was higher after blocked than interleaved encoding, 84.0% [83.0%, 84.8%] versus 79.2% [78.2%, 80.2%], bootstrap p <.001 (Fig. 3E). In contrast, interleaved sampling increased accessibility of the associated element B, with exact B identification increasing from 63.1% [61.3%, 64.9%] to 66.8% [64.9%, 68.6%], bootstrap p =.009 (Fig. 3F).

To determine whether this reflected relational binding rather than a general increase in B accessibility, we tested retrieval of the correct B against an equally familiar B lure. Correct-B forced-choice accuracy was greater following interleaved than blocked encoding, increasing from 92.4% [91.7%, 93.1%] to 93.9% [93.2%, 94.6%], bootstrap p <.001 (Fig. 3G). Permuting A-B assignments returned performance to the 50% baseline, all permutation p <.001, confirming that A cues preferentially retrieved their particular B associates.

We next counterbalanced whether A or B appeared first to distinguish interleaving from serial position. The correct-B versus lure similarity margin was greater following interleaved encoding in both orders, increasing by 0.0137 [0.0124, 0.0149] when A appeared first and by 0.0144 [0.0128, 0.0160] when B appeared first, both bootstrap p <.001 (Fig. 3H). The schedule-by-order interaction was close to zero, 0.0007 [-0.0013, 0.0027], p =.503, indicating that the effect did not depend on which element appeared first.

To test the transition account more directly, we replaced the binary blocked-versus-interleaved contrast with a graded manipulation of transition frequency. Each sequence contained the same three samples from A and B, but their temporal ordering produced one, three, or five transitions. Correct-B forced-choice accuracy increased from 92.4% [91.7%, 93.1%] with one transition to 93.4% [92.7%, 94.0%] with three and 93.9% [93.1%, 94.6%] with five transitions (Fig. 3I), both bootstrap p <.001. Thus, relational retrieval strengthened progressively as sampling transitioned more frequently between associated elements.

Lastly, we directly tested whether the mnemonic effect of a transition depended on preceding sampling history. We compared identical A3-B2 transitions at the same serial position following A1-A2-B1-**A3-B2**-B3 versus A1-B1-A2-**A3-B2**-B3. Both sequences contained the same visual samples and total number of between-element transitions. Before the critical transition, correct-associate accuracy was higher following ABAA than AABA sampling, 89.07% [87.75%, 90.32%] versus 85.88% [84.37%, 87.28%], bootstrap p <.001. After the identical transition, accuracy increased to 94.78% [93.70%, 95.80%] and 93.33% [92.17%, 94.42%], respectively. Critically, the initially weaker AABA history showed a greater improvement, yielding a significant history-by-transition interaction, gain difference = 1.73% [0.80, 2.67], p =.004 (Fig. 3J). Thus, the mnemonic contribution of an identical between-element transition depended on the representational state established by preceding samples, beyond transition frequency or exposure alone.

Together, these findings extend the sequential-sampling account from individual-element representations to relational memory. Relational retrieval strengthened as sampling moved more frequently between associated elements, while the contribution of an individual transition depended on the sampling history that preceded it. The temporal organisation of sampling may therefore provide a mechanism through which separately sampled elements become integrated into a relational memory representation.

### Sequential sampling transforms the geometry of element representations

Having shown that sequential sampling shapes both the individuation of element representations and relations between elements, we next examined how the geometry of an individual-element representation changes as successive samples are integrated.

As additional fixations were integrated, the population representation became progressively lower-dimensional (Fig. 4A). The normalised participation ratio, which measures how broadly variance is distributed across representational dimensions, decreased from 0.156 in the first two-fixation window (95% CI [0.153, 0.160]) to 0.109 in the final window [0.106, 0.112], bootstrap p <.001. Thus, variation across element representations became increasingly concentrated along fewer dominant dimensions as evidence accumulated. Importantly, this reduction in population dimensionality does not imply that individual states contained less information. Semantic category information was already strongly represented early in encoding, with class-balanced accuracy of 0.881 [0.868, 0.894] in the first window and 0.861 [0.836, 0.886] in the final window. By contrast, exact-image access increased from 0.709 [0.694, 0.726] to 0.903 [0.888, 0.919], bootstrap p <.001 (Fig. 4B). Thus, declining dimensionality did not reflect progressive abstraction away from element-specific information. Semantic information was available early, whereas exact-element information continued to increase.

**Figure 4.**
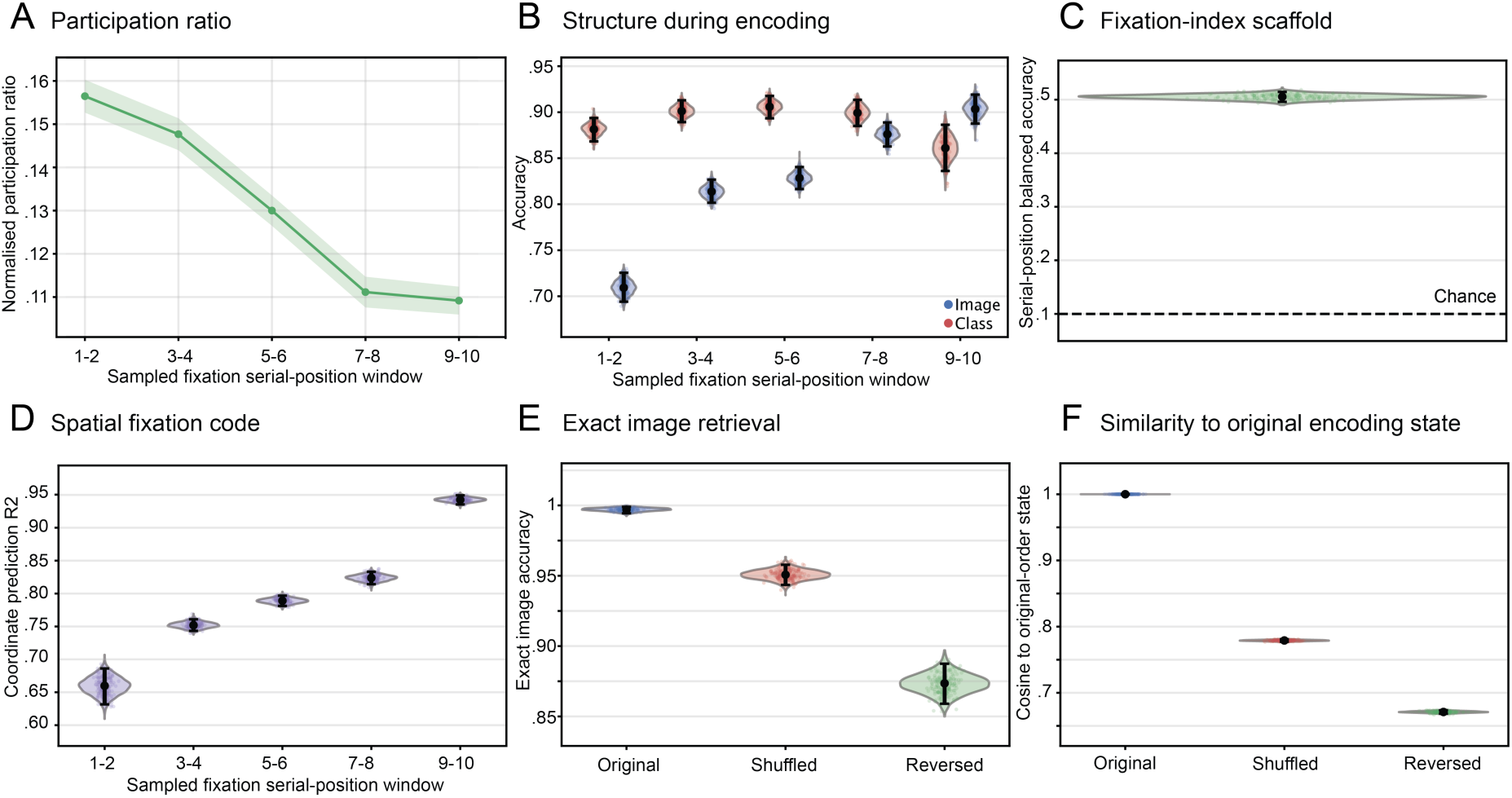
Sequential sampling progressively transforms latent representations into retrieval-ready memory states. **A** Effective dimensionality of the population representation decreased as successive samples were integrated. Normalised participation ratio is shown across serial-position windows, indicating that variance across recurrent states became increasingly concentrated along fewer dominant dimensions. B Semantic class information (red) was strongly decodable from early recurrent states, whereas exact-image access (blue) increased as additional fixations were integrated. C A linear probe trained and evaluated using image-grouped cross-validation decoded fixation serial position substantially above chance, indicating that the recurrent state preserved the temporal structure of sampling. D Image-grouped ridge regression recovered fixation coordinates from the recurrent state, with spatial decoding increasing across encoding. E Fixation order contributed to exact-image retrieval: presenting the same fixation tokens and coordinates in shuffled or reversed order reduced access to the corresponding stored image. F Order perturbation also changed the latent representation, with shuffled and reversed sequences producing final recurrent states that were less similar to the original-order state.For A-D, uncertainty was estimated using paired, class-stratified bootstrap resampling of images. For E-F, shuffled-order estimates were first averaged across deterministic permutations within each image and then evaluated using the same paired bootstrap procedure. Violin plots show bootstrap distributions, faint points show sampled bootstrap estimates, and black markers and error bars denote the observed estimate and bootstrap 95% confidence interval. Bootstrap repetitions quantify image-sampling uncertainty and do not represent independent agents or independently trained models.

The recurrent state also retained information about how it had been sampled. Fixation serial position was decoded with an accuracy of 0.505 [0.496, 0.514], substantially above the 0.10 chance level (Fig. 4C). The x-y coordinates of the current fixation were also recoverable throughout encoding, with coordinate-decoding performance increasing from *R*^2^ (coefficient of determination) = 0.660 [0.632, 0.686] in the first window to *R*^2^ = 0.943 [0.935, 0.949] in the final window, bootstrap p <.001 (Fig. 4D). Because fixation position and serial position are partly coupled by the ordered trajectory, this establishes coordinate decodability rather than spatial coding independent of temporal position.

We next asked whether this temporal structure merely remained decodable or actually affected memory. We presented exactly the same fixation tokens and coordinates in their original sequence, shuffled, or reversed, thereby isolating the contribution of temporal order. Changing fixation order selectively impaired retrieval of the specific element (Fig. 4E). Exact-image accuracy decreased from 0.997 [0.994, 0.999] in the original condition to 0.951 [0.943, 0.958] after shuffling and 0.873 [0.859, 0.887] after reversal, both bootstrap p <.001. Class retrieval remained near ceiling, decreasing only from 1.000 to 0.998 and 0.994, respectively. The order in which information was sampled therefore mattered much more for recovering the individual element than its semantic category.

The same manipulation altered the representation itself (Fig. 4F). Similarity to the original-order latent state decreased from 1.000 to 0.779 [0.777, 0.781] after shuffling and 0.671 [0.668, 0.674] after reversal, both bootstrap p <.001. The recurrent network therefore did not simply accumulate an unordered collection of glimpses: the sequence in which they occurred helped determine the resulting representation.

Together, these findings show that sequential sampling progressively reorganises individual-element representations along fewer dominant dimensions while preserving element-specific and sampling-history information. Crucially, temporal order contributes to the representation itself and specifically to distinguishing the individual element.

### Retrieval benefits from reinstating the original gaze trajectory

If sequential gaze sampling contributes to constructing the representations available to episodic memory, then retrieval should depend on how well the cue reinstates the way an element was originally sampled. To test this directly, test images were encoded using five-fixation trajectories, while five-fixation retrieval cues shared between one and five samples with the encoding trajectory (Fig. 5A). Exact-image retrieval increased monotonically with overlap, from 0.711 (95% bootstrap CI [0.691, 0.731]) at Intersection over Union (IoU) = 0.11 to 0.963 [0.955, 0.972] at complete overlap, bootstrap p <.001. Semantic-class retrieval was comparatively robust, increasing more modestly from 0.958 [0.949, 0.966] to 0.996 [0.993, 0.999], bootstrap p <.001. The same graded dependence was evident before the learned projector. In raw GRU state space, exact-image accuracy increased from 0.168 [0.152, 0.184] at IoU = 0.11 to 0.999 [0.997, 1.000] at complete overlap, bootstrap p <.001. Thus, even when cue length was held constant, reinstating more of the original sampling trajectory improved access to the specific stored element, and this effect was already present in the recurrent representation.

**Figure 5.**
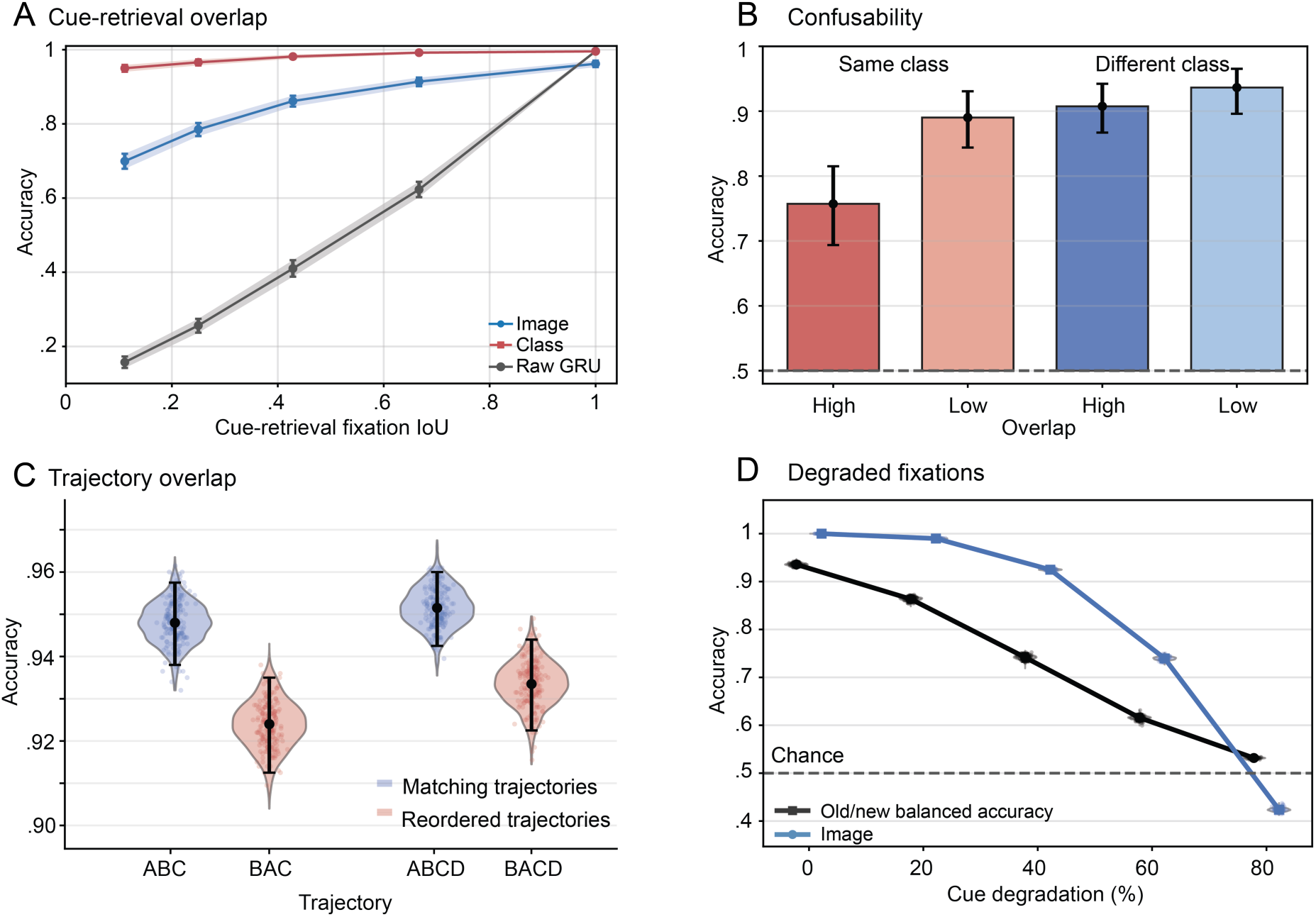
Memory retrieval depends on reinstatement of the original trajectory. **A** Exact-image retrieval increased strongly with loll in both the projected memory space (blue) and the raw GRU state space (grey), whereas projected semantic-class retrieval (red) remained comparatively robust at low overlap. Lines show observed accuracy; shaded regions and error bars show paired-bootstrap 95% confidence intervals. B Forced-choice retrieval of the correct associated B was lowest when the target A and competing A’ belonged to the same semantic class and had highly overlapping encoding trajectories. Bars and black points show observed accuracy, error bars show paired class-stratified bootstrap 95% confidence intervals, and the dashed line indicates 50% chance performance. **C** Exact-image retrieval was higher when retrieval cues preserved the encoding order than when the first two fixation samples were exchanged. Three-fixation sequences compared A-B-C with B-A-C, and four-fixation sequences compared A-B-C-D with B-A-C-D, holding sampled information, fixation locations, and the final sample(s) constant. Blue indicates matching order and red indicates swapped order. Violins and scattered points show paired-bootstrap estimates; black markers and error bars indicate observed accuracy and 95% bootstrap confidence intervals. D Progressive degradation of fixation cues preferentially impaired exact episodic retrieval. Exact-image retrieval (blue) as well as old/new recogntion (black) declined markedly as increasing proportions of fixation tokens were removed. Violins show hierarchical-bootstrap distributions, lines and markers show observed estimates, and error bars show 95% bootstrap confidence intervals. The dashed line indicates the 50% chance level for old/new recognition.

Reinstatement benefits alone are compatible with established encoding-specificity effects. We therefore tested the stronger prediction that sampling history should alter the similarity structure of memories themselves, determining which competing associations are retrieved. We constructed target-lure pairs in which A and a competing A’ each retrieved different arbitrary associates, B and B’ (Fig. 5B). Target and lure varied independently in semantic similarity and sampling-history overlap. Retrieval of the correct associate was lowest when A’ matched A in both semantic category and sampling history. Correct-B forced-choice accuracy was 0.757 for same-class, high-overlap lures, compared with 0.890 for same-class, low-overlap lures, 0.908 for different-class, high-overlap lures, and 0.936 for different-class, low-overlap lures. Accuracy in the same-class, high-overlap condition was lower than in each other condition, all bootstrap p <.001. Correct-B-versus-B’ similarity margins showed the same pattern, ranging from 0.379 for same-class, high-overlap lures to 0.696 for different-class, low-overlap lures. Thus, semantic similarity and sampling-history overlap jointly increased not only confusion between memory cues, but also retrieval of information from a stored competing associate.

To directly test history dependence at retrieval, we encoded each image using a sequence of 3 or 4 fixation samples and tested retrieval after exchanging the first two samples, leaving all sampled locations and remaining sample positions unchanged (Fig. 5C). Retrieval cues therefore contained identical visual samples and fixation locations, but the two first samples swapped position. Exact-image retrieval was higher for matching than swapped sequences: 94.8% [93.8, 95.75] versus 92.4% [91.25, 93.50] for three-fixation sequences and 95.15% [94.25, 96.00] versus 93.35% [92.25, 94.40] for four-fixation sequences, both bootstrap p <.001. Retrieval therefore depended on preceding sampling history even when the same information was available and later cue samples matched encoding.

Finally, we asked whether progressive degradation of fixation-derived cues could reproduce a behavioural phenomenon observed in humans. Wynn and colleagues (2020) parametrically degraded retrieval cues in an old/new paradigm to test how much visual evidence is required for pattern completion. We similarly replaced increasing proportions of each sampled fixation trajectory with an uninformative token (Fig. 5D). Exact-image retrieval declined from 1.000 [1.000, 1.000] with complete cues to 0.424 [0.417, 0.431] at 80% degradation, bootstrap p <.001, while semantic information remained more robust, with class accuracy of 0.570 [0.563, 0.576] under the most degraded condition. In an old/new recognition analogue, balanced accuracy likewise declined from 0.936 [0.930, 0.941] with complete cues to 0.532 [0.529, 0.535] at 80% degradation, bootstrap p <.001. Progressive degradation of fixation-derived cues therefore impaired both element-specific retrieval and recognition, qualitatively reproducing the behavioural effect reported by Wynn and colleagues ^64^.

Together, these findings suggest that sampling history contributes to the structure of the stored representation, making reinstatement of the original trajectory particularly important for recovering a specific element.

### Exploratory generative readout and correspondence with human neural activity

Our final analyses explored two extensions beyond the core computational account: the reconstructive consequences of fixation-constrained encoding and correspondence between model-derived representational dynamics and human neural activity. These analyses were exploratory and were not required to produce the preceding memory effects.

We first asked how an incomplete, fixation-constrained memory representation might support reconstruction of a more complete perceptual element. To test this, we trained a representation-conditional diffusion model (RCDM) using the projected final recurrent state together with a mask indicating which regions had been sampled during encoding. Reconstruction accuracy was strongly spatially selective (Fig. 6A-B), with mean squared error inside the fixation mask (0.070 ± 0.077 SD) being significantly lower than outside it (0.285 ± 0.209 SD, p <.001). The same pattern was observed for absolute reconstruction error. L1 error was lower inside the fixation mask (0.198 ± 0.118 SD) as compared to outside it (0.398 ± 0.169 SD, p <.001). The reconstructed images illustrated the same principle qualitatively (Fig. 6B). Sampled regions tended to preserve the corresponding image content, whereas unsampled regions showed more substitutions, omissions, and generative completion. Thus, when reconstruction was explicitly prioritised at locations that had been sampled during encoding, those regions could be recovered substantially more faithfully than regions that had not been sampled, which relyed more heavily on learned regularities

**Figure 6.**
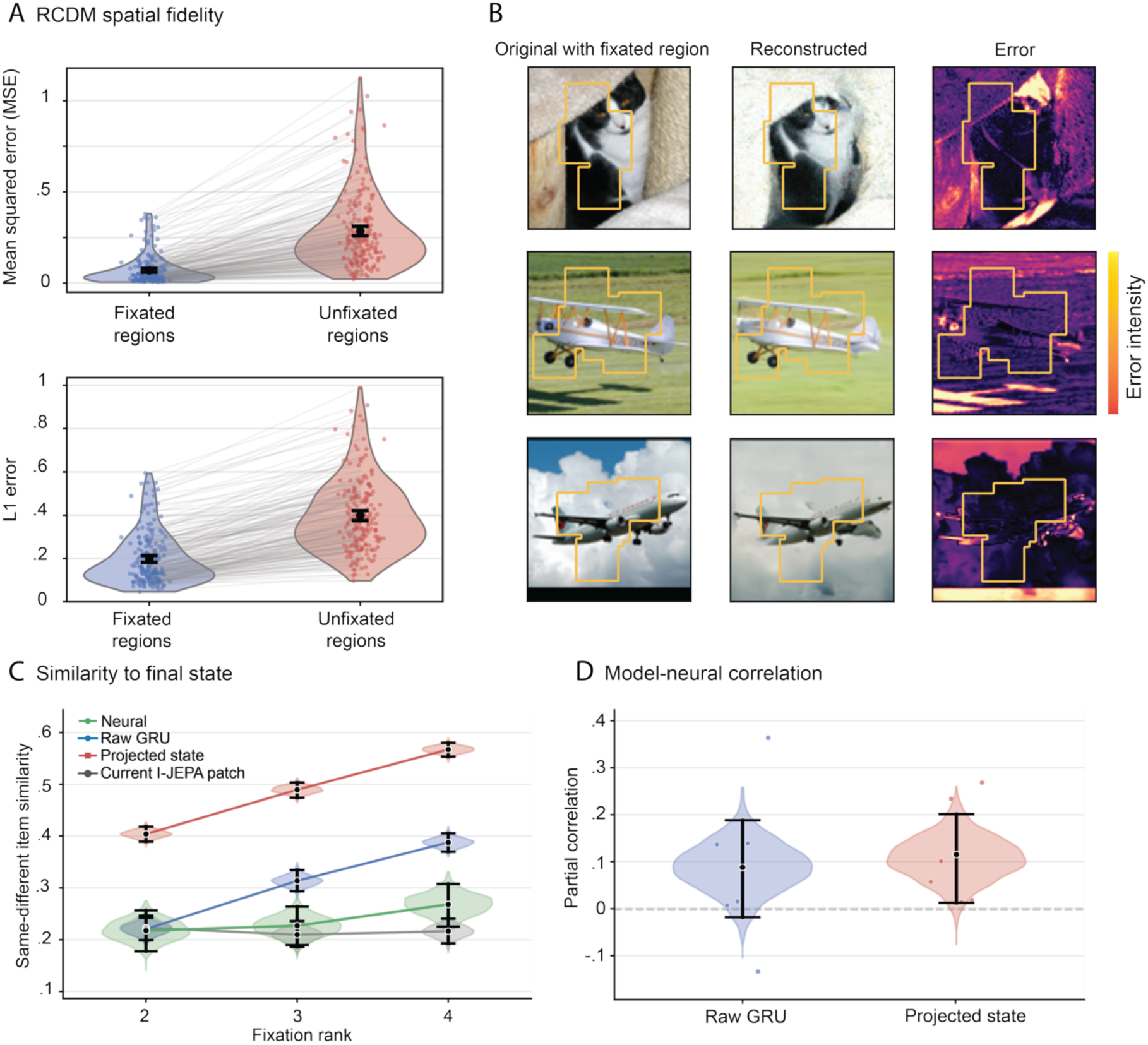
Exploratory generative readout and correspondence with human neural activity. **A** Reconstruction fidelity was compared between regions that had been directly sampled during encoding and regions that had not been fixated. When conditioned on the compressed episodic memory representation together with the fixation mask, the generative decoder reconstructed attended regions with substantially higher fidelity than unattended regions. Mean squared error was approximately fourfold lower within fixated regions than outside the fixation mask, with a similar effect observed for L1 reconstruction error. Individual points represent reconstructed images, grey lines connect paired measurements from the same image, and violin plots illustrate the full distribution. B Representative reconstructions illustrating fixation-dependent readout. For each example, the original image (left) is shown alongside the sampled fixation regions used during encoding (highlighted with yellow outline), the reconstructed image generated from the compressed episodic memory representation, and the corresponding pixel-wise reconstruction error. Reconstructions preserve substantially greater detail within attended regions, whereas unsampled portions of the scene rely more heavily on generative completion. C Neural and model representational measured as similarity across fixation ranks 2-4 to the same image’s fixation-7 representation minus similarity to other images. Neural activity from lateral temporal cortex is shown in green, raw GRU states in blue, projected states in red, and current l-JEPA patches in grey. Neural patterns were measured 100-300 ms after fixation onset following overlap correction. D Partial correlations between neural and model convergence after controlling for current-patch and running-average similarity, fixation rank, and timing/gaze variables. Coloured points represent individual participants. In C-D, violins show bootstrap distributions of equal-participant estimates; black markers and error bars indicate observed estimates and 95% bootstrap confidence intervals. Intervals quantify trial-sampling uncertainty within six fixed participants. The dashed line in D indicates zero correlation.

We next explored whether the model’s internal representations correspond to neural dynamics during natural viewing. Reanalysing data from Kragel and colleagues ^69^, in which they used simultaneous eye tracking and intracranial recordings from six epilepsy patients, we compared fixation-related activity patterns in lateral temporal cortex with representations generated by the frozen model from participants’ observed sampling trajectories. Neural patterns were measured 100-300 ms after fixation onset (correcting for overlapping responses; see Methods). At fixation ranks 2-4, convergence was quantified as similarity to the same image’s fixation-7 representation minus similarity to fixation-7 representations from other images in the same recording block. Neural and recurrent representations showed increasing similarity towards their respective final state, whereas the I-JEPA patch representations showed little increase (Fig. 6C). Projected-state similarity was positively associated with neural similarity after accounting for current-patch and running-average similarity, fixation rank, and timing/gaze variables (mean partial *r* =.115, 95% bootstrap CI [.012,.201], bootstrap p =.026; Fig. 6D). The corresponding association for the raw GRU state remained uncertain *r* =.088, [−.018,.188], bootstrap p =.104).

Together, these exploratory analyses illustrate potential consequences and empirical relevance of fixation-constrained representations. Sampled information provided stronger constraints on generative reconstruction, while projected-state convergence showed a positive association with neural representational convergence beyond the specified visual and timing controls.

## Discussion

Computational theories of episodic memory, including hippocampal indexing ^43,70,71^, Complementary Learning Systems ^38,72^, attractor-network models ^73,74^, and more recent generative accounts ^41,42^ have provided important insights into the potential mechanisms of how memories are stored, retrieved, and consolidated. However, these accounts typically begin from the assumption that a representation suitable for memory already exists. Here, we sought to extend these accounts by specifying how such representations can be constructed from sequential perceptual sampling before they are stored, integrating sampling into the memory computation itself. Fixation-dependent visual features and sampling locations are sequentially integrated by a recurrent network, projected into memory space, and stored in associative memory. At retrieval, partial sampling generates states through the same pathway, providing cues to the stored representation. Each fixation therefore updates a recurrent state already shaped by preceding samples, such that visual content, sampling location, and temporal history jointly determine the representation available for storage. Within an element, successive samples progressively individuate its representation, whereas transitions between elements promote relational integration. Crucially, this mechanism generalised to novel held-out images encountered once, after the recurrent and projection networks had been frozen. The architecture therefore separates slow learning of how sequential samples are transformed into memory representations from the one-shot construction and storage of individual episodic representations. Sampling history is not only contextual information attached to a memory, but helps determine the representation that is stored.

Within individual elements, successive samples progressively reduced uncertainty about identity (Fig. 2). Semantic category became distinguishable from competing categories before exact element identity, consistent with a broader temporal hierarchy in which gist can become available before finer detail ^56,57,59^. This is also consistent with evidence that information about individual objects is maintained and updated across eye movements, with successive fixation-derived features integrated into increasingly stable object representations ^75–79^. Established accounts propose that cortical object representations provide input to hippocampal episodic encoding ^80,81^. Our model adds a specific mechanism for constructing such input. The same visual sample can have different mnemonic consequences depending on the content, locations, and order of preceding samples. Successive samples increased separation of an element from same-category competitors without implementing hippocampal pattern separation. Because hippocampal pattern separation is thought to reduce representational overlap between similar experiences ^60^, we propose that sequential sampling may shape the similarity structure of the input on which this mechanism operates. This is consistent with evidence that visual sampling during encoding predicts later mnemonic discrimination of similar stimuli ^82–84^ and predicts that the evolving differentiation of the representation, rather than sampling amount alone, should determine mnemonic discrimination.

Improved memory with additional samples is not sufficient to distinguish our model from evidence accumulation accounts, which predict that more observations should improve memory ^35^. If accumulated observations are the same, their order should not change the final representation. We tested this directly by holding sampled visual information and fixation locations constant while changing only temporal order. Rearranging the same observations changed the resulting state and selectively impaired exact-image retrieval while largely preserving semantic-category retrieval (Fig. 4). Thus, the representation depends not only on which observations are accumulated, but on the trajectory through which they are encountered. Sequential integration also reorganised representational geometry: population dimensionality declined while element-specific information increased, even as sampling location and serial position remained decodable. This dimensionality transformation is consistent with theoretical accounts in episodic memory ^28^ and learning-related transformations in cortical and hippocampal population activity ^61–63^. Sequence sensitivity is not unique to our account, however. Temporal-context models explain order effects through a gradually changing internal context ^44–46^. Our model extends this idea by proposing that sampling history transforms the evolving element representation itself, rather than only associating an already defined element with an evolving context. It therefore predicts that fixation-by-fixation changes in the evolving representation should explain subsequent memory beyond temporal position and similarity between encoding and retrieval contexts.

Across elements, relational accessibility increased as sampling transitioned more frequently between A and B (Fig. 3). Holding sampled information and total exposure constant, this effect was graded across sequences containing one, three and five A-B transitions, suggesting that temporal organisation helps determine how strongly separately sampled elements become related in memory. Recent naturalistic evidence supports this account ^8^ where gaze preferentially transitioned between related scene objects during encoding, and stronger relation-guided scanning predicted later object and relational recall. Because this relationship was correlational, however, it could not determine whether relational transitions construct the memory or instead reflect successful scene understanding. Related work shows that temporal transition structure can shape representations acquired from sequential experience ^85^ and that different study curricula can emphasise different aspects of learned structure ^86^. Our model makes the more specific prediction that relational binding depends not only on transition frequency, but also on the sampling history preceding each transition. We tested this by presenting an identical A-B transition at the same serial position following differently ordered histories, while holding perceptual information, cumulative exposure, transition count and the immediately preceding sample constant (Fig. 3J). The same transition produced different gains in relational accessibility depending on preceding sampling history. Thus, its mnemonic contribution depends on the evolving representation into which information is integrated, supporting relational memory as a history-dependent construction rather than simply an accumulation of associations.

Sampling history also shaped retrieval (Fig. 5). Gaze during retrieval partially reinstates spatial and temporal sampling patterns established during encoding ^14,87,88^, and partial recovery of encoding-related sampling information has been associated with successful memory. In our model, cue length was held constant while overlap with the encoding trajectory varied. Greater overlap improved exact-image retrieval, whereas semantic-category retrieval depended less on overlap (Fig. 5A). This is consistent with encoding specificity ^89^, but goes further by showing that sampling history contributed to constructing the stored representation itself. Critically, when identical fixation samples were presented in a different order, exact-element retrieval was impaired even though the final sample, or final two samples, remained unchanged (Fig. 5C). Retrieval therefore operates on an evolving representation shaped by preceding cue samples, rather than simply accumulating matching perceptual information independently of temporal organisation.

This mechanism provides a possible link between gaze reinstatement and hippocampal pattern completion, which allows a stored representation to be recovered from a partial or degraded cue ^90^. Wynn et al. (2020) found that degraded retrieval cues elicited gaze patterns associated with previously encoded images, while neuroimaging provides a complementary link ^23,24,91^: hippocampal activity patterns associated with successful memory formation overlap with patterns predicting subsequent gaze reinstatement ^92^. In the model, partial reinstatement increased similarity between the retrieval cue and the stored representation. We therefore hypothesise that gaze reinstatement can contribute to hippocampal pattern completion by reinstating sampling-dependent components of an encoded representation. Beyond facilitating target retrieval, sampling history also influenced which memories competed. Retrieval was most disrupted when a competitor shared both semantic category and sampling history with the target (Fig. 5B) and this competition propagated to retrieval of associated information. This extends behavioural findings showing that gaze reinstatement can promote retrieval of a similar studied image from an incomplete lure ^64^ and predicts that hippocampal pattern completion should depend not only on perceptual similarity but also on sampling-history similarity. Consistent with this link, neural memory reactivation has been reported to align temporally with correctly reinstated gaze shifts ^93^.

Our results also suggest a possible division of computational roles between cortical and hippocampal systems. In the model, successive fixation-dependent samples are integrated in a recurrent state before the resulting representation is stored in associative memory, which does not feed back into the evolving state during encoding. This architecture is compatible with proposals that information is temporarily maintained and accumulated across saccades within transsaccadic working memory ^34–36,94,95^. Fixation-dependent information could therefore be progressively integrated across cortical visual and association networks; consistent with this possibility, recurrent prediction of successive visual glimpses produces integrated scene representations that increasingly align with human mid-and high-level visual cortex ^52^. The resulting sampling-dependent representation could then provide input to hippocampal associative-memory mechanisms, shaping the similarity structure on which pattern separation operates ^60^ and providing the cortical input available for relational binding. During retrieval, partial reinstatement of a sampling trajectory could provide a cortical cue for hippocampal pattern completion, followed by broader cortical reinstatement.

This feedforward separation is a simplification. Cortical and hippocampal systems can interact continuously during visual sampling. For example, Kragel and colleagues ^69^ identified short-term retrieval episodes during initial scene encoding, reflected in revisitation fixations that predicted later reinstatement and were preceded by hippocampal theta dynamics and increased top-down influence from hippocampus to the dorsal attention network ^92,96^. Sequential sampling may therefore form a recurrent loop in which perceptual sampling updates memory and memory guides subsequent sampling ^21,23,91^. The present model describes the first phase of this loop. Extending it with memory-guided sampling is an important direction for future work.

The exploratory generative analysis illustrates a further possible consequence of fixation-constrained encoding (Fig. 6A-B). Sampled regions were reconstructed more faithfully than unsampled regions, consistent with reconstructive accounts in which episode-specific information constrains reconstruction while poorly specified information is filled in using learned regularities ^97–99^. Because the decoder was explicitly trained to prioritise sampled regions, this should be interpreted as a prediction of fixation-prioritised reconstruction rather than an emergent property of the model. Supplementary analyses further showed that sequentially constructed representations can support downstream consolidation and semanticisation processes described by established memory models ^38,41,100^, but these processes are secondary to the central mechanism examined here. The exploratory neural analysis additionally revealed correspondence between the model’s internal representation and fixation-related activity patterns in lateral temporal cortex (Fig. 6C-D).

In sum, sequential sampling provides a candidate computational process through which representations available to episodic memory are constructed. Sampling within an element progressively individuates its representation, whereas transitions between elements strengthen relational integration. Because each update depends on the representation produced by preceding samples, sampling history becomes embedded in the resulting memory representation and shapes retrieval through trajectory reinstatement and competition between related memories. Episodic memory may therefore depend not only on what is experienced, but on the trajectory through which its elements are sampled and related over time.

## Acknowledgements

This research was funded by the Max Planck Society (C.K., S.D., E.C., C.D.). A.N. was supported by Marcus and Amelia Wallenberg Foundation (MAW2022.0048). The funders had no role in study design, data collection and analysis, decision to publish or preparation of the manuscript.

## Author contributions

The model was conceived and developed by C.K, S.D. and E.C. with consultation from A.N. and C.D. The manuscript was written by C.K. with input from D.D, E.C., A.N. and C.D.

## Data and Code availability

Data and code for reproducing the figures, running the analyses and retraining the model is available at Zenodo (https://doi.org/10.5281/zenodo.23206339).

## Methods

### Model Overview

The model implements element-specific memory as a sequence of gaze sampling (Fig. 1A), recurrent evidence accumulation, associative storage, and reconstructive readout. Visual inputs were represented using a pretrained Image-based Joint-Embedding Predictive Architecture (I-JEPA) model ^101^. For each STL-10 image, fixation coordinates selected the nearest I-JEPA patch token, converting the image into an ordered sequence of attended visual samples. Rather than encoding the full image at once, each memory was formed from a variable fixation trajectory containing between five and ten sampled fixations.

Sequential fixation input was integrated by a GRU-based GlimpseSequence recurrent neural network (RNN) ^48,50,102^. At each fixation, the RNN received both the attended I-JEPA patch token **z**_t_ and the corresponding fixation coordinates ***p****_t_* = (*x_t_*, *y_t_*),

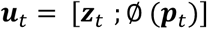

and updated its latent state according to:

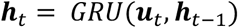

For each image, fixation trajectories were constructed by retaining the first fixation and then randomly sampling between four and nine additional fixations (yielding sequences of 5-10 fixations) from the recorded scanpath, with a centre-biased sampling distribution. The resulting variable fixation sequence therefore defined the gaze trajectory from which the episodic memory was constructed.

The evolving hidden state therefore acted as a recurrent evidence accumulator, progressively integrating the visual content, spatial sampling location, and temporal order of successive fixations (what, where, and when). The RNN was trained using a self-supervised next-token prediction objective,

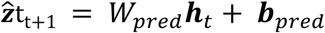

encouraging the latent state to predict the embedding of the next sampled fixation and thereby capture the structure of sequential visual exploration. The RNN was optimised by minimising the mean squared error between predicted and observed next-token embeddings across all fixation transitions,

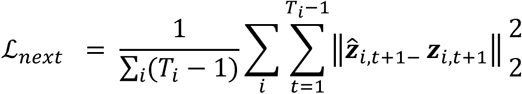

After training, the RNN parameters were frozen. A separate projection network mapped recurrent states into associative-memory space,

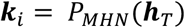

where *P_MHN_* consisted of a LayerNorm followed by a two-layer multilayer perceptron (Linear-GELU-Linear). The projection network was trained while keeping the RNN fixed, aligning projected partial-sequence states with the projected final state of the corresponding training image while maintaining discriminability from competing training images.

The pretrained I-JEPA encoder was kept fixed throughout, while the recurrent and projection networks were trained only on the STL-10 training split. For all primary memory analyses, all three components were frozen and applied to held-out STL-10 test images. Held-out STL-10 test images were then encoded once through the frozen pathway, and a new associative-memory bank was constructed by storing one projected final state for each test image as an MHN memory key. No learned parameters were updated during memory formation or retrieval. For analyses comparing all serial positions, each held-out image was encoded using the complete ten-fixation trajectory, ensuring that the same 2000 images contributed to every fixation position ^39^.

The core memory model therefore consisted of I-JEPA fixation representations, the GlimpseSequenceRNN, the projection network, and the MHN.

Retrieval was performed by passing a partial fixation sequence through the same frozen RNN and projection network to obtain a query state,

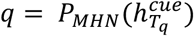

Retrieval scores were computed from the cosine similarity between the normalised query and each stored memory key,

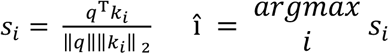

For the primary memory analyses, we used a held-out one-shot procedure with the STL-10 test split. The pretrained I-JEPA encoder was kept fixed throughout, and all learned parameters of the GlimpseSequenceRNN and projection network were frozen before memory formation. A new MHN memory bank was constructed from the held-out test images by encoding each image once and storing its final projected state as a new memory key. Intermediate fixation prefixes from the same held-out images were subsequently used as retrieval queries against this fresh test-image bank. No recurrent or projection parameters were updated during memory formation or retrieval. For analyses comparing all serial positions, each held-out image was encoded using the complete ten-fixation trajectory, ensuring that the same 2000 images contributed to every fixation position.

Additional decoders were used only as readout tools. A variational autoencoder (VAE; Kingma & Welling, 2013) was used to examine whether replayed or lesioned memory states reconstructed images with greater prototype structure and reduced element-specificity. A representation conditional diffusion model (RCDM; ^103^) was used as an exploratory decoder to test whether fixated regions could be reconstructed with higher spatial fidelity than non-fixated regions. These decoders were not part of the associative memory mechanism itself, but instead probed what information was preserved within the sequentially constructed RNN-MHN memory states.

To assess robustness across random initialisation, we repeated model training and the core analyses across five independent seeds. We quantified seed-level effects for uncertainty reduction, target-evidence gain, participation-ratio reduction, reinstatement, order sensitivity, lure confusability, and relational transitions (Supplemental Fig. 3).

### Statistics

Unless otherwise specified, inferential statistics reported for the primary analyses were obtained from a single fixed trained model and therefore quantify sampling uncertainty across images, associations and, where explicitly simulated for behavioural analogues, evaluation or task profiles, rather than variability across independently trained model instances. Bootstrap repetitions were not treated as independent model instances or repeated training runs.

### Visual input and dataset

The model was evaluated on the STL-10 dataset using its standard training and test splits. Images were processed by a pretrained I-JEPA vision transformer to obtain global image embeddings and spatially resolved patch-token representations. Each fixation coordinate was mapped to the nearest I-JEPA image patch, converting a spatial sampling trajectory into an ordered sequence of attended visual tokens.

Image identifiers and semantic class labels were retained throughout the analyses. The pretrained I-JEPA encoder was kept fixed throughout, while the GRU and projection network were trained using only the STL-10 training split. For all primary memory analyses, the GRU and projection network were subsequently frozen and held-out STL-10 test images were encoded once to form a new MHN memory bank, with one final projected representation stored for each image. Exact-image retrieval was defined relative to this newly constructed test-image bank. Thus, the primary analyses assessed whether the fixed architecture could construct and retrieve individual memories for novel images after a single encoding encounter, without updating any learned model parameters.

### Sequential sampling

Visual input was represented using patch-level features extracted from the pretrained I-JEPA model. Each fixation coordinate was assigned to its nearest I-JEPA image patch, producing an ordered sequence of attended patch tokens for each image (Fig. 1). The patch tokens and corresponding spatial coordinates jointly formed the sequential input to the memory model. Each complete sampling trajectory contained a maximum of ten fixations.

For the primary analyses, each image was encoded using a trajectory containing between five and ten fixations. The initial fixation was always retained, and the remaining fixations were sampled without replacement using a centre-biased weighting scheme. Selected fixations were subsequently presented in their original temporal order. This procedure produced a distinct variable-length encoding trajectory for each stored image.

For analyses comparing representations across all serial positions, all ten available fixation positions were instead retained for every held-out test image. This ensured that each serial position was estimated from the same set of 2000 images, rather than from progressively smaller subsets of variable-length trajectories.

Thus, individual-element representations were not derived from complete image representations presented at a single time point. Instead, they were progressively constructed from sequences of spatially localised visual samples. Sequential sampling therefore provided the perceptual input through which evidence was accumulated to construct the representation subsequently available to associative memory.

### Sequential evidence accumulation

Sequential samples were integrated using a gated recurrent unit (GRU), implemented as a GlimpseSequenceRNN. At each fixation t, the network received the corresponding I-JEPA patch token together with an embedding of its spatial coordinates. The GRU updated its hidden state recurrently, such that ℎ*_t_* represented the visual and spatial evidence accumulated across all fixations observed up to that point. The final hidden state provided an integrated representation of the sampled element.

The GRU had a hidden dimension of 768. Fixation coordinates were transformed by a multilayer perceptron into a 32-dimensional spatial embedding before being combined with the visual patch representation. The recurrent network was trained using a self-supervised next-token prediction objective. Given the samples observed up to fixation t, the network predicted the I-JEPA representation of the patch attended at fixation *t*_+1_. Training minimised the mean squared error between predicted and observed next-patch representations:

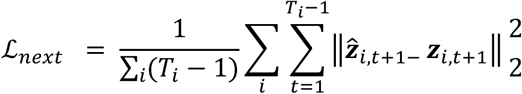

Where ***z****_i_*_,*t*+1_ denotes the observed I-JEPA patch representation and ***ẑ***<*_i,t+1_* its prediction from the preceding recurrent state. This objective encouraged the hidden state to retain information about the sequentially sampled visual input that was predictive of subsequent attended content. The recurrent network was trained only on images from the STL-10 training split for 200 epochs with a learning rate of 2 ∗ 10^−4^ and a batch size of 128.

### Projection and associative memory

After training the fixation-sequence model, the GRU parameters were frozen and a projection head was trained to map recurrent states into associative-memory space. The projection head comprised LayerNorm, linear, GELU, and linear transformations. Its intermediate dimension was 1024 and its output dimension was 768, matching the GRU hidden-state dimensionality. The projector was trained to transform states generated from partial fixation sequences into representations approximating the final state of the same sampled trajectory.

During training, a variable-length fixation trajectory was sampled for each image. Prefixes containing one, two, three, or five fixations, when available, were processed by the frozen GRU and projection head. Each partial-sequence representation was aligned with the projected final state generated from the same sampled trajectory for that image. The objective combined retrieval cross-entropy across images within each batch, cosine alignment, and mean squared error between partial and final representations. Retrieval logits were calculated from cosine similarities using an inverse temperature of 10. The projection head was trained for 200 epochs with a learning rate of 1 ∗ 10^−4^.

During projector training, representations from the STL-10 training split supplied the learning signal. The trained projector was subsequently frozen. For the primary memory analyses, held-out STL-10 test images were encoded once through the frozen I-JEPA, GRU, and projection pathway, and their projected final states were stored as new keys in a freshly constructed Modern Hopfield Network (MHN). No training-set memories were included in these banks and no learned parameters were updated during memory formation or retrieval. Memory keys therefore reflected representations constructed through sequential sampling rather than global I-JEPA image embeddings.

Retrieval followed the same recurrent and projection pathway used to construct the memory keys. A partial or perturbed fixation sequence was processed by the frozen GRU and projector to generate an MHN query. Queries and stored keys were L2-normalised, and retrieval scores were computed from their cosine similarity. For autoassociative element retrieval, stored memories were ranked by query-key similarity, with the highest-scoring key defining top-1 retrieval. Thus, exact-image retrieval measured retrieval of newly stored held-out elements rather than recognition of images encountered during GRU or projector training. Heteroassociative retrieval used in the relational analyses is described separately below.

### Linear decoding of representational content

To determine what information was accessible from recurrent states generated for previously unseen images, we performed held-out state-code analyses using 2000 STL-10 test images. The pretrained I-JEPA encoder and trained GRU were frozen; no parameters of the visual encoder, recurrent network, projector, or associative-memory system were updated. Each test image was processed using its complete ten-fixation trajectory, ensuring that the same images contributed to every serial position. Linear probes were used solely as post hoc readouts of information present in the recurrent representation.

For linear decoding, each 768-dimensional recurrent state was transformed using a fixed random Gaussian projection to 128 dimensions. Semantic information was assessed within five two-fixation windows: fixations 1-2, 3-4, 5-6, 7-8, and 9-10. At each fixation position, a ridge classifier predicted the STL-10 class from the projected recurrent state. Performance was evaluated with stratified 10-fold cross-validation. All observations from a given image were assigned to the same fold, preventing states from that image from appearing in both probe-training and probe-test data. Predictions were pooled across held-out folds, and performance was quantified as class-balanced accuracy.

Image-specific access was assessed directly from the unprojected recurrent states. Each intermediate state was L2-normalised and compared by cosine similarity with the bank of final recurrent states for all held-out test images. Exact-image top-1 accuracy indicated whether the most similar final state belonged to the same image as the intermediate query. Measurements from adjacent positions were pooled into the same five two-fixation windows used for class decoding. This measure involved no fitted readout.

Temporal information was assessed with a ridge classifier trained to predict the serial position of each recurrent state within the ten-fixation sequence. Cross-validation was grouped by image, such that all observations from a given image remained within one fold. Pooled held-out predictions were evaluated using balanced accuracy across the ten possible fixation positions; chance performance was 0.10.

Spatial information was assessed using ridge regression from the projected recurrent state to the x-y coordinates of the current fixation. Coordinate predictions were generated out of fold using the same image-grouped 10-fold partitioning. Performance was quantified using the coefficient of determination, ***R*^2^**, calculated from pooled held-out predictions and averaged within the five two-fixation windows.

Uncertainty was estimated using 2000 paired, class-stratified bootstrap repetitions over held-out images. Images were sampled with replacement within each STL-10 class, with the same resampled images and multiplicities retained across fixation windows. All observations from a sampled image were retained together. We report observed out-of-fold estimates and percentile-based 95% bootstrap confidence intervals. Bootstrap repetitions quantify image-sampling uncertainty conditional on one fixed trained model, rather than independent agents or retrained model instances.

### Relational retrieval confusability

To test whether semantic similarity and gaze history jointly increased interference with associative retrieval, we performed a held-out target-lure analysis using the frozen I-JEPA encoder, GRU, and projection pathway. Each held-out test image was encoded once from a newly sampled variable fixation trajectory, and its final projected state was used as an A-key representation. Each A image was assigned a distinct arbitrary B associate using a fixed random one-to-one permutation of the held-out image set. Thus, a target A and competing lure A′ were paired with separate values, B and B′.

For each target image, a retrieval cue was generated from the first three fixations of its own encoding trajectory. Candidate A′ lures were selected by their proximity to the target in global I-JEPA image-embedding space and assigned to one of four conditions: same-class with high fixation overlap, same-class with low fixation overlap, different-class with high fixation overlap, or different-class with low fixation overlap. Fixation overlap was calculated between the complete sampled encoding trajectories of A and A′. High overlap required at least five shared fixation indices, whereas low overlap required no more than one shared index.

For each target-lure comparison, the target A and lure A′ representations were stored as the two keys of a heteroassociative MHN, with B and B′ stored as their corresponding values. The partial A cue queried this two-association memory, producing a soft retrieved value representation. Retrieval was evaluated by comparing the retrieved value with B and B′. Correct-associated retrieval was scored when the retrieved state was more similar to B than to B′, yielding a two-alternative chance level of 0.50. The relational margin was defined as the similarity to B minus the similarity to B′.

To reduce ceiling performance, each I-JEPA fixation token in the retrieval cue was independently corrupted with cosine-calibrated noise targeting a cosine similarity of 0.30 to its uncorrupted token. Fixation coordinates, stored A keys, and B values were unchanged. This analysis therefore tested whether a competing cue representation sharing both semantic category and sampling history with A preferentially caused retrieval of its associated value B′.

Statistical uncertainty was estimated using 2000 paired, class-stratified bootstrap repetitions over target images. The same resampled targets were retained across all four conditions, and primary contrasts were restricted to targets with valid lures in every condition.

### Uncertainty reduction during sequential sampling

To characterise how sequential sampling progressively constructed element-specific representations, we quantified retrieval uncertainty after each successive fixation. For this analysis, 2000 held-out STL-10 test images were encoded once through the frozen model and their final projected states stored in a newly constructed associative-memory bank. Every available prefix of each image’s stored fixation trajectory was then processed by the frozen GRU and projection head to generate an intermediate query. The query was compared with all stored memory keys using cosine similarity, and similarities were transformed into retrieval probabilities using a softmax function with an inverse temperature of 10 (Direction of evidence accumulation was preserved across β = 5, 10, and 20 and temperature-independent similarity margins showed identical increases across conditions).

Uncertainty was evaluated at both the image and semantic-class levels. Image-level entropy was calculated over the probability distribution across individual memory keys. Class probabilities were obtained by summing the probabilities of all stored memories belonging to each STL-10 category, from which class-level entropy was calculated. Entropy values were divided by their theoretical maxima,

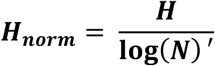

where N denotes the number of possible outcomes ***N_memories_*** for image entropy and ***N_classes_*** for class entropy, placing both measures on a common scale from 0 to 1.

For each fixation prefix, we additionally measured the probability assigned to the target image, the probability assigned to its semantic class, the rank of the target key, top-1 retrieval accuracy, and the margin separating the target from its strongest competing memory. Fixation-wise changes were calculated as the difference between successive prefixes, allowing the rate of uncertainty reduction and evidence accumulation to be examined across the encoding trajectory. Mean trajectories and fixation-wise changes were evaluated using 2000 paired, class-stratified image bootstraps, with identical resampled images retained across fixation prefixes. Early-to-late contrasts were restricted to images with observations at both prefixes.

These analyses quantified how additional fixation samples altered the distribution of evidence across competing memories and allowed the emergence of semantic category information to be compared with the development of image-specific episodic evidence.

To compare the emergence of semantic and element-specific information without relying on arbitrary absolute-probability thresholds, we performed a separate readout analysis in the raw GRU state space, before projection into associative-memory space. For the same 2000 held-out test images, each intermediate GRU state was compared with the bank of final GRU states from those images. Cosine similarities were transformed into probabilities using the same softmax procedure. The class margin was defined as the probability assigned to the correct class minus that assigned to the strongest competing class. The exact-image margin was defined as the probability assigned to the target image minus that assigned to the strongest competing memory.

For each image, stable discriminability was defined as the first fixation prefix at which the corresponding margin became positive and remained positive throughout all subsequent available prefixes. A positive class margin indicated that the correct category received stronger evidence than every competing category, whereas a positive image margin indicated that the exact target received stronger evidence than every competing memory. Uncertainty was estimated using 2000 paired, class-stratified image bootstraps. Images were resampled with replacement within STL-10 classes, with the same resampled images retained across fixation prefixes and across class-and image-level measures. Crossing times were calculated within each bootstrap repetition, and the class-image timing difference was evaluated from its paired bootstrap distribution. Results are reported as observed means with percentile 95% bootstrap confidence intervals.

### Encoding-retrieval fixation overlap

To test whether retrieval depended on reinstating the encoding trajectory without relying on unequal naturally occurring overlap levels, we conducted a held-out one-shot-memory analysis. The GRU and state projector were trained on the STL-10 training set and then frozen. We used 2000 novel STL-10 test images, none of which had contributed to training the GRU or projector. For each image, a new episodic memory was formed by processing a variable encoding trajectory through the frozen GRU and projector and storing its final projected state as a key in a fresh MHN. Thus, test images were added as memory contents, but no trainable model parameters were updated.

Encoding trajectories contained five fixation indices sampled without replacement from the ten available positions using the same centre-biased weighting as variable-fixation encoding, with the first fixation always retained. Retrieval trajectories also contained five fixations. For each image, retrieval trajectories were constructed to share exactly *m* = *1, 2, 3, 4*, or *5* fixation indices with its encoding trajectory. The first fixation was shared in every condition; the remaining shared indices were sampled from the encoding trajectory, and non-shared indices were sampled from positions not used during encoding. All selected indices were returned to their original temporal order.

Fixation overlap was quantified using intersection over union,

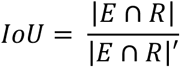

where ***E*** and ***R*** denote the encoding and retrieval fixation-index sets. Because both trajectories contained five fixation indices, the controlled overlap conditions corresponded to IoU values of 0.11, 0.25, 0.43, 0.67, and 1.00. Every held-out image contributed one observation to each overlap condition, equating image identity, sample size, and trajectory length across conditions.

Retrieval cues were corrupted with cosine-calibrated token noise with a target cosine similarity of 0.65, processed by the frozen GRU and state projector, and queried against the fresh MHN using the same one-step MHN update and inverse temperature used elsewhere. Exact-image top-1 accuracy was defined as retrieval of the correct held-out memory key; semantic class top-1 accuracy was defined as retrieval of a key with the correct STL-10 class.

To determine whether the effect depended on the learned projector, we repeated the analysis directly in raw GRU state space. Final raw GRU states from encoding trajectories were stored in a separate fresh MHN, and raw GRU states generated from the controlled retrieval trajectories were used as queries. This control used the same images, trajectories, noise, MHN retrieval procedure, and evaluation measures; only the learned projection from GRU state to memory space was bypassed.

Uncertainty was estimated with 2000 paired bootstrap repetitions over held-out images. The same resampled images and multiplicities were retained across all overlap conditions, preserving the paired structure of the manipulation. We report observed accuracies, percentile-based 95% bootstrap confidence intervals, and two-sided bootstrap probabilities for paired contrasts relative to the one-shared-fixation condition. Bootstrap repetitions quantify uncertainty from held-out image sampling and were not treated as independent agents, model instances, or training runs.

### Encoding-retrieval order reinstatement

To test whether retrieval depended on temporal order independently of fixation-set overlap, we conducted a held-out one-shot-memory analysis using 2000 STL-10 test images. The trained GRU and state projector remained frozen throughout. Each image was encoded using either its first three fixation samples, A-B-C, or its first four samples, A-B-C-D. Letters denote fixation-token–coordinate pairs from the same image. Final projected encoding states were stored as keys in separate fresh MHN banks for the three-and four-fixation conditions, with one key per image.

Retrieval cues contained the same samples as encoding, presented either in their original order or with the first two token-coordinate pairs exchanged: B-A-C or B-A-C-D. Sampled visual information, fixation locations, sequence length, and fixation-set overlap were therefore identical across order conditions, with IoU fixed at 1. The final fixation remained unchanged in three-sample sequences, and the final two fixations remained unchanged in four-sample sequences.

Retrieval tokens were corrupted with cosine-calibrated noise targeting a similarity of 0.65 to the uncorrupted token. Each sample was corrupted once before reordering, ensuring that identical corrupted token-coordinate pairs were used across matching and swapped conditions. The same noise realisations were also retained across sequence lengths and representational spaces. Encoding inputs were uncorrupted.

Retrieval sequences were processed by the frozen GRU and projector to generate queries to the corresponding MHN bank. Retrieval used one MHN update with an inverse temperature of 20. Exact-image top-1 accuracy indicated whether the highest-weighted memory key represented the queried image.

Uncertainty was estimated using 2000 paired, class-stratified bootstrap repetitions over held-out images. Images were resampled with replacement within STL-10 classes, with identical resampled indices and multiplicities retained across sequence lengths, order conditions, and representational spaces. Paired contrasts quantified swapped-minus-matching retrieval performance. We report observed estimates, percentile-based 95% bootstrap confidence intervals, and two-sided bootstrap probabilities. Bootstrap repetitions quantified image-sampling uncertainty from a fixed trained model and were not treated as independent agents or training runs.

### Dimensionality of recurrent representations

Effective dimensionality of the recurrent population representation was quantified using the participation ratio. For each fixation position, the 768-dimensional GRU states were collected across images and mean-centred. We then calculated the sample covariance matrix

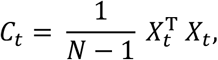

Where *X_t_* contains the centred recurrent states at fixation position *t*. Given the eigenvalues ℷ_j_ of *C_t_*, participation ratio was defined as,

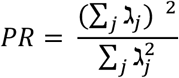

Participation ratio estimates the number of dimensions over which variance is effectively distributed. Larger values indicate that variance across images is distributed over more representational dimensions, whereas smaller values indicate that variance is concentrated within a lower-dimensional subspace. This is a population-level measure of representational geometry and does not imply that individual state vectors became smaller or necessarily contained less information.

PR was divided by the GRU hidden dimensionality of 768 to obtain a normalised participation ratio. 2000 STL-10 test images were evaluated with the pretrained I-JEPA encoder and trained GRU held fixed. Each image was processed using the complete ten-fixation trajectory, ensuring that the same images contributed to every fixation position. PR was calculated separately at each position and then averaged within five non-overlapping windows: fixations 1-2, 3-4, 5-6, 7-8, and 9-10.

Uncertainty was estimated using 2000 paired, class-stratified image bootstrap repetitions. Images were resampled with replacement within STL-10 classes, and the same resampled images were retained across all fixation positions and windows. For each repetition, covariance matrices and participation ratios were recalculated from the resampled recurrent states. Because duplicate image resamples can bias a covariance-based participation ratio downward, bootstrap repetitions were used to estimate the standard error and 95% confidence intervals were centred on the observed statistic using 1.96 times this standard error. The change in effective dimensionality was evaluated from the paired bootstrap distribution of the difference between the first and final fixation windows.

### Order-perturbation analysis

To determine whether retrieval depended on the temporal organisation of sampling for novel episodes, we performed a held-out one-shot order-perturbation analysis without retraining the I-JEPA encoder, recurrent network, or state projector. We used 2000 STL-10 test images, each encoded with the complete ten-fixation trajectory. A separate MHN bank was constructed by storing the projected final state generated from the original-order trajectory of each held-out image.

In the original-order condition, fixation tokens and coordinates were presented in their original temporal order. In the shuffled-order condition, the same token-coordinate pairs were randomly permuted within each image. In the reversed-order condition, the same sequence was presented in reverse. Thus, sampled visual information, fixation locations, and the set of ten fixation indices were identical across conditions; only their temporal order differed.

Each sequence was processed by the frozen RNN to produce a final hidden state, which was transformed into a query by the trained state projector. Queries were compared by cosine similarity with the held-out MHN keys generated from the original-order trajectories. Exact-image retrieval was scored as correct when the highest-ranked stored key represented the target test image. Class retrieval was scored as correct when the highest-ranked key belonged to the target STL-10 category. State preservation was quantified as the cosine similarity between the final RNN state generated by each perturbed sequence and the corresponding original-order state.

To account for variability among possible random permutations, the shuffled condition was evaluated using ten deterministic random permutations per image. Per-image outcomes were averaged across these permutations before statistical inference. The original and reversed conditions were each evaluated once because their temporal ordering was uniquely defined.

To match the degraded retrieval regime used in the confusability analysis, projected queries were corrupted with cosine-calibrated noise targeting cosine similarity of 0.30 relative to the uncorrupted query. The same base noise realisation was used across order conditions and shuffle repetitions, ensuring that condition differences reflected temporal order rather than different noise samples.

Statistical uncertainty was estimated using 2000 paired, class-stratified bootstrap repetitions over held-out images. Images were sampled with replacement within each STL-10 class, and the same resampled images and multiplicities were used across all order conditions. Paired contrasts quantified change from original to shuffled and from original to reversed order. We report observed estimates, percentile-based 95% bootstrap confidence intervals, and bootstrap probabilities for paired contrasts. Bootstrap repetitions quantify image-sampling uncertainty conditional on one fixed trained model, rather than independent agents or retrained model instances.

### Retrieval under degraded cue information

To examine how retrieval depended on the amount of information available at test, we progressively degraded sampled fixation trajectories used to query episodic memory. This analysis was inspired by degraded-cue recognition paradigms, but degradation was applied to sequential samples rather than image pixels. The held-out STL-10 test set was divided into balanced, disjoint sets of 1000 images. One set formed a newly constructed one-shot MHN memory bank, with one projected final state stored for each image; the other set was never stored and served as novel lures in the old/new recognition analogue. The pretrained I-JEPA encoder, GRU, and projection head remained frozen throughout.

Each held-out image was assigned an image-specific, centre-biased trajectory containing five to ten fixation samples, with the first fixation retained during trajectory construction. The final state from the complete trajectory formed the stored MHN key for old images. At retrieval, the full variable-length trajectory was used as the cue. At each degradation level, each valid fixation token was independently retained with probability equal to the visible-token fraction; non-retained tokens were replaced by a mean token calculated across the old and novel held-out sets. Fixation coordinates, temporal positions, and sequence length were retained. Padded positions outside the sampled trajectory were excluded, and at least one valid token was retained whenever the visible fraction was greater than zero. Thus, degradation removed local visual evidence while preserving the spatial and temporal scaffold of the sampled trajectory.

Degradation was evaluated at 0%, 20%, 40%, 60%, and 80%, corresponding to expected visible-token fractions of 1.0, 0.8, 0.6, 0.4, and 0.2. Retained tokens were additionally corrupted with cosine-calibrated noise before processing by the frozen GRU and projection head. For each evaluation profile, the target cosine was generated by adding normally distributed variation (SD = 0.05) to a nominal value of 0.90 and restricting the result to the range 0.45-0.85. In the reported run, effective profile-specific target cosines ranged from 0.784 to 0.850 (mean = 0.846). Each resulting query was compared with all 1000 held-out MHN keys. Exact-image accuracy indicated whether the highest-ranked key represented the queried stored image, whereas class accuracy indicated whether it belonged to the correct STL-10 category.

To model variability in evaluation conditions without retraining the network, we generated 50 participant-like evaluation profiles. Profiles differed in their random cue-degradation and noise draws, profile-specific target cosine, and old/new recognition criterion. All profiles evaluated the complete held-out old and novel sets, although image order was independently randomised within profile. Criterion bias was sampled from a normal distribution with an SD of 0.015. Each profile’s item assignment, target cosine, and criterion bias were retained across degradation levels; only the stochastic token-retention and noise draws varied. These profiles represented variation in cue degradation and decision conditions, not independently trained model instantiations.

An old/new recognition analogue was derived from the maximum cosine similarity between each query and the held-out MHN memory bank. Stored held-out images served as old items, whereas the disjoint held-out set served as novel lures. A common baseline criterion was defined as the midpoint between the pooled mean oldness scores for old and novel items under the 0% degradation condition. Each profile’s criterion bias was added to this baseline. Queries whose maximum similarity equalled or exceeded the resulting criterion were classified as old. Recognition was summarised using hit rate, false-alarm rate, correct-rejection rate, balanced accuracy, sensitivity (*d’*), and the separation between old and novel similarity distributions.

Uncertainty was estimated using 2000 paired hierarchical-bootstrap repetitions. Evaluation profiles were resampled with replacement, followed by resampling old and novel image observations within each selected profile. The same resampled observations were retained across degradation levels, preserving the paired structure of the manipulation. Observed estimates are reported with percentile 95% bootstrap confidence intervals, and differences from the complete-cue condition were evaluated from the corresponding paired bootstrap distributions.

### Fixation-constrained image association

To examine whether fixation-constrained representations could support associations between stimuli, we conducted an exploratory held-out one-shot A-B associative-memory analysis. The I-JEPA encoder, GRU, and projection head were trained on the STL-10 training split and then frozen. For each analysis, novel images from the STL-10 test split were encoded into fresh memory representations without any further parameter updates. Fifty task-sampling profiles each contained five independently sampled study blocks of 24 arbitrary A-B image pairs. Pair assignments were generated randomly, so semantic similarity did not define the association.

For each image, a centre-biased trajectory of five fixation samples was selected and processed by the frozen GRU and projection head to form its final projected representation. Within each study block, final A representations were stored as keys and the corresponding final B representations as values in a heteroassociative MHN. Retrieval was evaluated only among the 24 A-B associations in the current block; chance top-1 associative accuracy was therefore 1/24 (4.17%).

At retrieval, prefixes containing one to five sampled A fixations were processed through the same frozen GRU and projection head. The I-JEPA fixation-token embeddings in each cue prefix were corrupted using cosine-calibrated noise with a target cosine similarity of 0.30; fixation coordinates were retained. The softly retrieved B-valued state was compared with the 24 candidate B representations in the block. Associative accuracy indicated whether the closest candidate was the B image paired with the cue A. Cosine similarities between the softly retrieved B-valued state and the 24 candidate B representations were transformed with a softmax with inverse temperature 10 to estimate the probability assigned to the correct associate.

To determine whether associative cueing depended specifically on regions sampled during encoding, individual fixation regions from A were evaluated separately. A-region diagnosticity was defined as the query similarity to the target A key minus similarity to the strongest same-class competing A key within the study block. Fixation regions included in A’s stored five-sample trajectory were greedily matched to unselected regions from the same image on this diagnosticity measure. The primary matched analysis retained pairs with an absolute diagnosticity difference no greater than 0.02, thereby controlling for how effectively a region identified A while testing whether participation in encoding conferred additional access to B.

Sampling uncertainty was estimated using 2000 paired hierarchical-bootstrap repetitions. Task-sampling profiles were resampled with replacement, followed by A-B pairs within each selected profile; all cue-prefix measurements from the same pair were retained together. For the sampled-versus-unsampled analysis, region-level matches were averaged within A-B pair before resampling, while preserving the paired observations. We report observed estimates, percentile 95% bootstrap confidence intervals, and two-sided bootstrap p-values. Bootstrap repetitions quantify task and element sampling uncertainty from a fixed trained model, not independent model instances.

### Temporal organisation of associative encoding

To test whether the temporal organisation of sampling influenced the information retained in a joint representation, we constructed episodes containing three fixation samples from each member of an A-B image pair. The same fixation tokens and coordinates were presented in either blocked order (A_1_, A_2_, A_3_, B_1_, B_2_, B_3_) or interspersed order (A_1_, B_1_, A_2_, B_2_, A_3_, B_3_). The pretrained I-JEPA encoder, GRU, and projection head remained frozen, and the GRU hidden state was not reset within an episode. The final projected state after all six inputs therefore represented the complete ordered A-B sequence. One such episode representation was stored for each of 24 pairs in a study block.

All A-B pairs were sampled from the held-out STL-10 test split. For each task profile, joint episode representations and the separate A and B comparison banks were constructed de novo from these images and stored once in a new MHN; the pretrained I-JEPA encoder, GRU, and projector were not updated during associative encoding or retrieval.

Retrieval was tested separately using A_1_, A_2_, and A_3_. Each fixation token was corrupted using cosine-calibrated noise with a target cosine similarity of 0.65, processed individually through the frozen GRU and projection head, and compared by cosine similarity with the 24 episode representations from the corresponding encoding condition. The same corrupted cues were used for blocked and interspersed retrieval. Retrieval outcomes were averaged across the three A cues within each pair.

The softly retrieved MHN state was compared separately with banks containing the final projected within-image representations of the 24 A images and 24 B images. A retrieval was correct when the nearest A representation belonged to the cue element, whereas B accessibility was correct when the nearest B representation belonged to its assigned associate. Similarities were additionally transformed using a softmax with an inverse temperature of 10 to estimate the probability assigned to the correct A and B images. Chance performance for A and B top-1 retrieval was 1/24, or 4.17%.

To determine whether differences in B accessibility reflected pair-specific relational binding rather than general B accessibility, the correct B was additionally tested against a familiar B lure from the same study block. For each target, the lure was matched as closely as possible on standalone accessibility from its own fixation cues and, where possible, was selected from the same semantic class. Forced-choice accuracy indicated whether the softly retrieved state was more similar to the correct B than to the matched lure. Relational margin was defined as the similarity to the correct B minus the similarity to the lure. The corresponding two-alternative target probability was obtained by applying a softmax to these two similarities.

To separate temporal interleaving from serial position, the blocked and interleaved schedules were repeated with element order counterbalanced. The four schedules were blocked A-first A_1_, A_2_, A_3_, B_1_, B_2_, B_3_, blocked B-first B_1_, B_2_, B_3_, A_1_, A_2_, A_3_, interleaved A-first A_1_, B_1_, A_2_, B_2_, A_3_, B_3_, and interleaved B-first B_1_, A_1_, B_2_, A_2_, B_3_, A_3_. Relational margins were compared across schedules to test whether the interleaving effect depended on which element occurred first or occupied the final serial position.

A complementary transition-dose analysis replaced the binary schedule comparison with a graded manipulation of temporal transitions. We generated every order-preserving interleaving of three A and three B samples containing exactly one, three, or five transitions between elements. Within-element order was retained in every sequence. The one-and five-transition conditions each contained two schedules, whereas the three-transition condition contained eight schedules. Outcomes were averaged across schedules within each transition count, thereby balancing whether A or B appeared first. Forced-choice accuracy, relational margin, target probability, and exact B accessibility were evaluated as functions of transition count. Linear slopes were estimated across one, three, and five transitions.

As an empirical relational-memory baseline, A-B assignments were randomly permuted within each study block while retaining the same cues, candidate B representations, schedules, and serial orders. Unlike a derangement, permutations were unrestricted and therefore retained the number of fixed points expected by chance. This produced empirical null distributions centred near 4.17% for 24-way B accessibility, 50% for forced-choice retrieval, and zero for relational margins.

The analysis comprised 50 task-sampling profiles, each containing five independently sampled blocks of 24 arbitrary A-B pairs. All trained parameters remained fixed across profiles. Sampling uncertainty was estimated using 2000 paired hierarchical-bootstrap repetitions, resampling profiles and A-B pairs within profiles while retaining all schedule and transition conditions together. Permutation null distributions were based on 2000 within-block A-B assignment permutations. We report observed estimates with percentile 95% bootstrap confidence intervals and two-sided bootstrap or permutation p-values.

To test whether the consequence of an identical between-element transition depended on preceding sampling history, we compared two sequences: A1-A2-B1-A3-B2-B3 and A1-B1-A2-A3-B2-B3. Both contained the same fixation tokens and coordinates, three samples from each element, and three between-element transitions. Their final three inputs were identical, placing the critical A3-B2 transition at positions 4-5 in both histories. The manipulation therefore changed the order of the first three samples while matching cumulative exposure and transition count at each evaluated position.

The frozen GRU integrated each sequence continuously. Projected states were extracted after positions 4 and 5 and stored in separate autoassociative MHN banks for each history and stage, with 24 episode representations per bank. These prefix banks assessed relational accessibility if the episode were stored at each stage. The same noisy single-fixation A1-A2-A3 cues queried every bank using one MHN update with an inverse temperature of 20. Retrieved states were compared with the standalone B representation bank. Similarity scores were averaged across the three cues within each pair before calculating correct-B forced-choice accuracy against the familiarity-matched lure and the corresponding relational margin. Cue noise and lure identity were held constant across histories and stages.

History effects were evaluated separately at each stage. Transition gain was defined as performance at position 5 minus performance at position 4, and the history-by-stage interaction compared this gain between the ABA and AAB histories. The analysis used 50 task profiles, each containing five blocks of 24 arbitrary held-out A-B pairs. Uncertainty was estimated using 2000 paired hierarchical-bootstrap repetitions, resampling profiles, blocks within profiles, and pairs within blocks while retaining all history and stage conditions together. We report percentile 95% confidence intervals and two-sided bootstrap probabilities. Within-block A-B assignment permutations provided empirical relational-memory baselines.

### Masked generative readout

To investigate how episodic memory representations could be transformed back into image-like representations, we developed an exploratory masked generative readout based on a representation-conditioned diffusion model (RCDM). This decoder was not part of the core episodic-memory model and did not contribute to sampling, recurrent encoding, or MHN retrieval. Instead, it served as a computational probe of whether a recurrent memory representation, together with explicit information about where visual input had been sampled, could support spatially selective reconstruction.

For each STL-10 training image, the global conditioning vector was derived from the projected final state of the sequential sampling model. This representation corresponded to the memory state used to construct the image’s associative-memory key. A binary spatial mask was generated from the fixation locations sampled during encoding. The mask was resized to the decoder resolution and concatenated with the noisy RGB image as an additional input channel to the diffusion U-Net at every denoising step. The decoder therefore received both a global representation of the accumulated fixation sequence and an explicit spatial representation of the sampled image regions.

The RCDM was trained using a conditional diffusion-denoising objective. At each training step, Gaussian noise was added to the target image, and the network learned to predict the corresponding denoising target conditioned jointly on the recurrent memory representation and fixation mask. To formalise the assumption that sampled regions provide higher-confidence episodic evidence than unsampled regions, the denoising loss was spatially weighted:

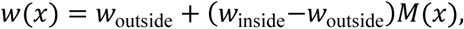

where *M*(*x*) in {0,1}denotes whether pixel *x* fell within the fixation mask. Pixels inside the mask received a weight of *w*_inside_ = 1.0, whereas pixels outside it received a weight of *w*_outside_ = 0.1. The weighted denoising objective was therefore

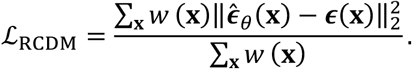

The decoder was still trained to reconstruct the entire image, but errors within sampled regions contributed more strongly to parameter updates. This weighting was introduced deliberately to test the computational consequences of fixation-prioritised reconstruction; spatially selective fidelity should therefore not be interpreted as an emergent property of an otherwise spatially uniform decoder.

The decoder was trained on all 5000 STL-10 training images for 437500 optimisation steps. Reconstructions were generated from the exponential-moving-average model checkpoint using DDIM sampling with 100 denoising steps.

Spatial reconstruction fidelity was evaluated across 200 generated images. For each image, MSE and L1 error relative to the original image were calculated separately inside and outside the fixation mask. The primary contrast was the paired outside-minus-inside error, with positive values indicating greater reconstruction error in unsampled regions.

Uncertainty was estimated using 2000 class-stratified paired-bootstrap repetitions. Images were resampled with replacement within STL-10 class, and identical resampled images were used to calculate inside-mask error, outside-mask error, and their paired difference. Observed estimates are reported with percentile 95% bootstrap confidence intervals and two-sided bootstrap p-values.

### Neural-model representational correlation

To explore correspondence between model-derived representations and human neural dynamics, we analysed simultaneous intracranial recordings and eye tracking from six participants in the publicly available Kragel et al. (2021) dataset ^69^. Analyses were restricted to encoding trials involving natural images from COCO and anatomically defined lateral temporal cortex (LTC). LTC selection followed the authors’ Harvard–Oxford atlas procedure. Bipolar channels qualified when either constituent contact belonged to LTC; hippocampal pairs were excluded. Eye events were aligned to intracranial recordings using native EyeLink timestamps, synchronisation markers and the authors’ trial-alignment rules.

Intracranial and eye-tracking data were preprocessed following Kragel et al. (2021), with adaptations for Python implementation. In short, data were bipolar-referenced, resampled to 500 Hz and corrected for line noise at 60, 120 and 180 Hz. Channel-specific interictal-discharge masks were derived using the released detection procedure. For the present analyses, data were further resampled to 100 Hz using a 101-tap anti-aliasing FIR filter. Artifact masks were expanded by 120 ms to accommodate filter support and a safety margin.

Each participant’s observed study scanpaths were processed through the frozen I-JEPA encoder, GRU and projection head. Fixation coordinates selected the nearest spatial I-JEPA patch token and were rescaled to the model’s coordinate frame. No model parameters were updated and no artificial cue noise was introduced.

Overlapping fixation-related responses were accounted for using a time-expanded regression-deconvolution model following the framework of ^104^, implemented in Python. The model included fixation and scene onset/offset events, fixation duration, preceding saccade size, fixation rank, gaze position and elapsed time, with response kernels spanning-300 to +800 ms. Single-fixation patterns retained the focal event’s fitted contribution and residual activity.

The first model-valid fixation was excluded because it frequently began before image onset, confounding its fixation-aligned response with scene onset. The GRU was reset before fixation 2. Fixations 2-7 were then processed continuously. Queries were evaluated at ranks 2, 3 and 4, using fixation 7 as a later reference rather than an independently established final memory state. Fixation 7 was used as a trade-off between enough trials and as late as possible in the encoding sequence. All fixations and their complete 100-300-ms response windows had to fall within the trial, with fixation onsets at least 200 ms after image onset. Fixation 7 onset additionally had to follow fixation-4 onset by at least 440 ms. First presentations satisfying these criteria and neural-quality requirements yielded 413 trials.

Neural patterns comprised mean corrected voltage during 100-300 ms after fixation onset, calculated separately for each bipolar channel and divided by that channel’s within-block voltage SD. Convergence was defined as cosine similarity to the same image’s fixation-7 pattern minus mean similarity to fixation-7 patterns from other images in the same block. Each same-versus-different comparison used identical channels across query ranks and reference conditions, requiring at least eight valid channels and three eligible comparison images. Equivalent contrasts were calculated for raw GRU states, projected states and current-patch tokens. Model comparisons used the same eligible reference images as the neural analysis; no MHN retrieval step was involved.

Model-neural correspondence was quantified separately for raw GRU and projected states using partial Pearson correlations. Neural convergence, model convergence and covariates were centred within trial to remove stable trial-level offsets. Ordinary least-squares residualisation accounted for categorical fixation rank, elapsed time since scene onset, time to the reference fixation, fixation duration, gaze x-y position, displacement between successive retained query positions, current-patch convergence and running-average patch convergence. Displacement was set to zero for the first retained query. Correlations were calculated between the residual neural and recurrent convergence measures.

Uncertainty was estimated using 2000 paired bootstrap repetitions, resampling whole trials with replacement within each participant and recording block while retaining all query ranks together. Nuisance regressions and correlations were recalculated in each repetition. Group estimates were equal-weight averages of participant estimates, with independently combined participant bootstrap repetitions. We report percentile 95% confidence intervals and descriptive two-sided bootstrap tail probabilities, calculated as twice the smaller zero-crossing tail proportion with a plus-one correction, capped at one. These estimates quantify trial-sampling uncertainty within six fixed participants, conditional on the reference banks and fitted deconvolution kernels; they do not estimate generalisation across participants. The analyses, including response-window and reference-rank selection, were exploratory and inferential summaries were unadjusted.

## Supplemental Material

### Methods

#### Memory decay

To examine how episodic-memory accessibility changed with memory age, we conducted a held-out one-shot decay analysis. The pretrained I-JEPA encoder, GRU, and projection head remained frozen. A fresh MHN was constructed from 2,000 STL-10 test images, each encoded once through a newly sampled variable trajectory of five to ten fixations. Projected final RNN states were inserted sequentially in 20 blocks of 100 images, with one key stored per image.

Newly stored memories were assigned an initial strength of 1.0. Before each subsequent block was inserted, strengths of all previously stored memories were multiplied by a decay factor of 0.97:

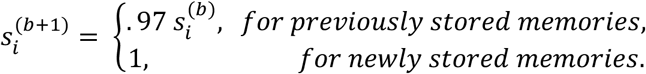

Decay altered the accessibility of stored keys without removing or modifying their representational content. Retrieval scores combined cosine similarity between the L2-normalised query and memory key with a logarithmic strength bias:

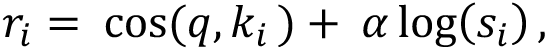

where *α* =1.0 controlled the influence of memory strength. Thus, older and weaker memories received a progressively larger retrieval penalty while remaining available in the memory bank.

Retrieval queries were generated from the first three fixation samples of each image’s own stored encoding trajectory. At each storage stage, up to 200 images were evaluated from every available memory block; because blocks contained 100 images, all stored images were evaluated. Exact-image retrieval indicated whether the highest-scoring key represented the queried image, whereas class retrieval indicated whether it belonged to the correct STL-10 category. Performance was summarised across all stored memories and separately for the oldest and most recently added blocks.

Uncertainty was estimated using 2000 class-stratified bootstrap repetitions. Within each storage stage and memory-age condition, images were resampled with replacement separately within semantic classes. Exact-image and class accuracy were calculated from the same resampled images, preserving their paired relationship. Observed estimates are reported with percentile 95% bootstrap confidence intervals. Memory-strength trajectories were deterministic consequences of the specified decay rule and were plotted without uncertainty intervals.

### Replay-driven cortical learning

To examine how replay selected by episodic memory could shape downstream cortical representations, we trained a separate convolutional variational autoencoder (VAE). The VAE was not involved in sampling, recurrent encoding, or associative retrieval. Instead, it served as an exploratory model of replay-driven cortical learning and reconstructive readout. The I-JEPA encoder, GRU, and projection head were trained on the STL-10 training split and then frozen. For this analysis, 2000 held-out STL-10 test images were encoded once into a fresh MHN using newly sampled variable trajectories of five to ten fixations.

Episodic memories were inserted into the MHN in 20 blocks of 100 images. Before each new block was added, the strengths of previously stored memories were multiplied by 0.97, whereas newly stored memories were assigned a strength of 1.0. Retrieval scores combined cosine similarity between the partial-cue query and stored key with the logarithmic strength bias described in the memory-decay analysis. Queries were generated from the first three fixation samples of each image’s own stored encoding trajectory.

After each insertion block, up to 1,000 partial-fixation cues queried the memories currently available in the MHN. The retrieved memory index selected the associated original test image for replay to the cortical VAE. Fewer replay events were available during the earliest blocks because fewer than 1000 memories had yet been stored. Cortical learning was therefore determined by episodes selected through strength-weighted MHN retrieval rather than uniform exposure to the held-out image set.

Following each replay cycle, the VAE was trained for 20 epochs with a learning rate of 1 ∗ 10^−4^. Its latent dimensionality was 512, the KL-divergence term was weighted by 0.05, and gradient norms were clipped at 1.0.

At evaluation, partial fixation cues first retrieved the highest-scoring memory through the MHN. The original image associated with that retrieved memory was then passed through the VAE encoder and decoder:

Partial cue - MHN retrieval - retrieved image - VAE reconstruction

This explicit associative-retrieval stage allowed successful access to a stored memory slot to be distinguished from the image-specific information preserved in its cortical reconstruction. Exact-image and class-level identification were assessed by matching reconstructed images to the corresponding held-out reference representations.

Representational geometry was evaluated across 400 images, comprising up to 20 images from each of the 20 storage blocks. Geometry was quantified in raw pixel space, an independent frozen full-image VQ-VAE feature space, and the latent space of the replay-trained VAE. Analyses measured within-class variance, distance from individual representations to their class prototype, and pairwise distances between class centroids. Compression was calculated as the corresponding value for the original representations minus that for the reconstructed representations. Positive values therefore indicated reduced within-class variability, movement towards class prototypes, or reduced between-class separation following reconstruction.

Uncertainty was estimated using 2,000 class-stratified bootstrap repetitions. Images were resampled with replacement within class, with identical resampled indices retained for original and reconstructed representations across all feature spaces. Class prototypes and between-class centroids were recomputed within each bootstrap sample. We report observed descriptive values with percentile 95% bootstrap confidence intervals and two-sided bootstrap p-values for compression contrasts.

### Memory lesion

To examine what information remained recoverable when associative episodic retrieval was unavailable, we evaluated a cortical-only retrieval pathway following the held-out replay-driven learning procedure described above. The I-JEPA encoder, GRU, and projection head remained frozen after training on the STL-10 training split. A fresh MHN was constructed from 2000 held-out test images, each encoded once through a newly sampled variable trajectory of five to ten fixations.

The cortical VAE and a linear cue-to-latent readout were trained throughout the same 20 blocks of MHN-mediated replay. After each block of 100 newly stored memories, up to 1000 replay events were selected through strength-weighted MHN retrieval. The images associated with retrieved memories were used to update the VAE. In parallel, the projected partial-cue representations were paired with the VAE latent representations of replay-selected images and used to train the linear cortical readout. The VAE and readout were therefore trained on the same episodes selected through associative replay.

Following each replay block, the linear readout was trained for four epochs with a learning rate of 1 ∗ 10^−P^ and a batch size of 128. Its objective was to predict the 512-dimensional VAE latent representation associated with a replayed image directly from the projected representation generated by the first three fixation samples. This established a cortical pathway that could map partial cues into the representational space learned through replay.

After all 2000 memories had been stored and replay-driven training was complete, the VAE and linear readout were frozen. The intact pathway used the partial fixation cue to select a stored episode through the MHN before reconstructing its associated image:

Partial cue - MHN retrieval - retrieved image - VAE reconstruction

For the lesion condition, access to MHN-mediated retrieval was removed by setting all memory strengths to zero and bypassing the MHN selection step. The same projected partial cue was instead mapped directly into VAE latent space by the frozen linear cortical readout and decoded without selecting a stored memory:

Partial cue - linear cortical readout - VAE latent - decoded image

The critical manipulation was therefore removal of associative selection of an individual stored episode, rather than degradation of the encoder, cortical VAE, cortical readout, or evaluation cues. The intact and cortical-only pathways differed only in whether the partial cue could access an individual stored episode through MHN-mediated retrieval.

Evaluation used the same 400 images sampled across the 20 storage blocks as in the replay analysis. Cortical-only reconstructions were assessed for exact-image and class-level identification. Representational geometry was quantified in pixel space, an independent frozen full-image VQ-VAE feature space, and the latent space of the replay-trained VAE. Analyses measured within-class variance, distance to class prototypes, and distances between class centroids. Positive compression values indicated that cortical-only reconstructions had lower variability or smaller representational distances than the corresponding original images.

Uncertainty was estimated using 2,000 class-stratified bootstrap repetitions. Images were resampled with replacement within class, with identical resampled indices retained for original and cortical-only representations. Prototypes and class centroids were recomputed within each bootstrap sample. Observed descriptive values are reported with percentile 95% bootstrap confidence intervals and two-sided bootstrap p-values for compression contrasts.

## Results

### Memory decay

As memory strength decayed across subsequent storage blocks, exact retrieval showed a pronounced recency gradient (Fig. 5A). For the same oldest memory block, exact-image accuracy declined from 0.990 (95% bootstrap CI [0.970, 1.000]) after block 5 to 0.000 after block 20, bootstrap p <.001. In contrast, the most recently stored block remained at 1.000 exact-image accuracy at the final stage.

Semantic retrieval was substantially more resistant to the same manipulation (Fig. 5B). For the oldest memories, class top-1 accuracy declined from 1.000 after block 5 to 0.900 [0.840, 0.950] after block 20, bootstrap p <.001 Thus, progressive weakening of associative-memory traces preferentially disrupted access to individual episodes while preserving much of their semantic category information.

### Memory degradation preferentially preserves semantic structure

Previous generative models have shown that replay-trained cortical representations become more prototypical and exhibit reduced variability within semantic categories ^41^. However, reduced within-class variability alone cannot distinguish selective abstraction towards meaningful category structure from a more general collapse of the representational space. We therefore characterised this transformation along three complementary dimensions: variation among exemplars within a class, distance of individual representations from their class prototype, and separation between different class centroids. This allowed us to ask whether memory degradation specifically removed element-level detail while preserving the broader organisation separating semantic categories.

We next examined the consequences of replaying held-out episodic memories into a downstream cortical model (Supplemental Fig. 2C). A fresh associative memory bank was constructed from 2000 test images while the I-JEPA encoder, GRU, and projection head remained frozen. Memories retrieved from this bank trained a separate cortical VAE; the VAE was exposed only to replay-selected images rather than uniformly to the held-out set. Associative retrieval continued to select the correct stored memory with high accuracy, with MHN exact-image retrieval reaching 0.985. However, exact-image identification after VAE reconstruction was only 0.098. Thus, associative memory still selected the appropriate episode, but its cortical reconstruction expressed substantially less image-specific information.

In pixel space, this loss of specificity took the form of selective movement towards semantic prototypes. Within-class variance decreased from 0.0619 [0.0579, 0.0628] for original images to 0.0373 [0.0343, 0.0384] after reconstruction. Distance to the corresponding class prototype decreased from 0.1201 [0.1117, 0.1219] to 0.0661 [0.0608, 0.0679]. Prototype compression was 0.0539 [0.0497, 0.0550], whereas between-class compression was only 0.0061 [0.0081, 0.0112]. Prototype compression therefore exceeded between-class compression, difference = 0.0479 [0.0403, 0.0452], bootstrap p <.001. Reconstructed images became more prototypical without a comparable collapse of the broader separation between semantic categories.

The independent frozen VQ-VAE showed the same directional pattern. Within-class variance decreased from 0.0114 [0.0102, 0.0120] to 0.0092 [0.0082, 0.0099], and prototype distance decreased from 0.2044 [0.1813, 0.2185] to 0.1620 [0.1428, 0.1745]. Prototype compression was 0.0423 [0.0364, 0.0458], compared with between-class compression of 0.0301 [0.0220, 0.0449]. This numerical difference was smaller than in pixel space and did not reliably differ from zero, difference = 0.0122 [-0.0063, 0.0212], bootstrap p =.210. Thus, the independent feature space supported reduced within-category variation and movement towards prototypes, although evidence for preferential prototype compression over between-class compression was not conclusive.

The replay-trained VAE’s own latent space behaved differently. Within-class variance decreased from 0.00181 to 0.00164, and prototype distance decreased from 0.7307 to 0.6037. However, between-class compression was 0.3667 [0.3086, 0.3632], substantially greater than prototype compression of 0.1270 [0.1037, 0.1222], difference =-0.2397 [-0.2493,-0.1975], bootstrap p <.001. The plastic VAE latent space therefore underwent broad global contraction, even though the decoded reconstructions showed more selective prototype-oriented change in pixel space.

Replay thus reduced image-specific information and promoted category-biased reconstruction, but the apparent geometry depended on the measurement space. The decoded visual outputs showed clear selective movement towards class prototypes in pixel space, whereas the internal latent space of the plastic cortical model showed broader contraction of representational geometry.

Finally, we asked what information remained when associative-memory retrieval was no longer available (Supplemental Fig. 2D). The replay-trained cortical VAE and linear cue-to-latent readout were trained using the same held-out memories selected through MHN-mediated replay and then frozen. In the intact pathway, a partial fixation cue selected an individual stored memory through the MHN before reconstruction. In the lesion condition, MHN-mediated selection was bypassed and the same cue was mapped directly into cortical latent space. This isolated information recoverable through the cortical pathway without access to an individual stored episode. Exact-image identification from cortical-only reconstructions was 0.000, whereas class-level identification was 0.115, indicating that the direct cortical pathway did not independently recover episodic identity and retained only weak category information.

The geometry of cortical-only reconstructions showed pronounced loss of image-specific structure. In pixel space, within-class variance decreased from 0.0619 [0.0579, 0.0628] to 0.0027 [0.0024, 0.0028], corresponding to compression of 0.0592 [0.0553, 0.0601], bootstrap p <.001. Prototype distance decreased from 0.1201 [0.1116, 0.1222] to 0.0045 [0.0041, 0.0046], bootstrap p <.001. Between-class compression was much smaller, 0.0115 [0.0136, 0.0235]. Prototype compression exceeded between-class compression by 0.1041 [0.0871, 0.1010], bootstrap p <.001. Cortical-only reconstructions therefore converged strongly towards class prototypes while preserving comparatively more separation between category centroids.

The frozen VQ-VAE showed the same selective pattern. Within-class variance decreased from 0.0114 [0.0101, 0.0120] to 0.0014 [0.0012, 0.0015], bootstrap p <.001. Prototype distance decreased from 0.2044 [0.1808, 0.2176] to 0.0218 [0.0187, 0.0238], bootstrap p <.001. Between-class compression was 0.0371 [0.0147, 0.0843], and prototype compression exceeded between-class compression by 0.1455 [0.0885, 0.1709], bootstrap p <.001. Thus, an independent visual feature space confirmed that bypassing associative retrieval preferentially removed distinctions among individual images within semantic categories.

The replay-trained VAE latent space again showed broad contraction. Within-class variance decreased from 0.00181 to 0.00041, while prototype distance decreased from 0.7307 to 0.1108. Prototype compression was 0.6199 [0.5693, 0.5963] and between-class compression was similarly large, 0.6208 [0.6063, 0.6713]. Thus, unlike pixel and frozen VQ-VAE spaces, the plastic VAE latent space did not show a meaningful selective advantage for prototype over between-class compression; both dimensions contracted strongly.

Together, these analyses reveal a dissociation between image-specific and semantic structure. Progressive weakening of associative-memory strength preferentially disrupted access to individual stored episodes while preserving category information. Replay and removal of associative retrieval shifted reconstructed visual outputs towards category prototypes and reduced distinctions among exemplars, while preserving relatively more of the broader organisation separating semantic classes. The plastic cortical VAE latent space was less selective, showing more general contraction.

---

**Supplemental Figure 1.**
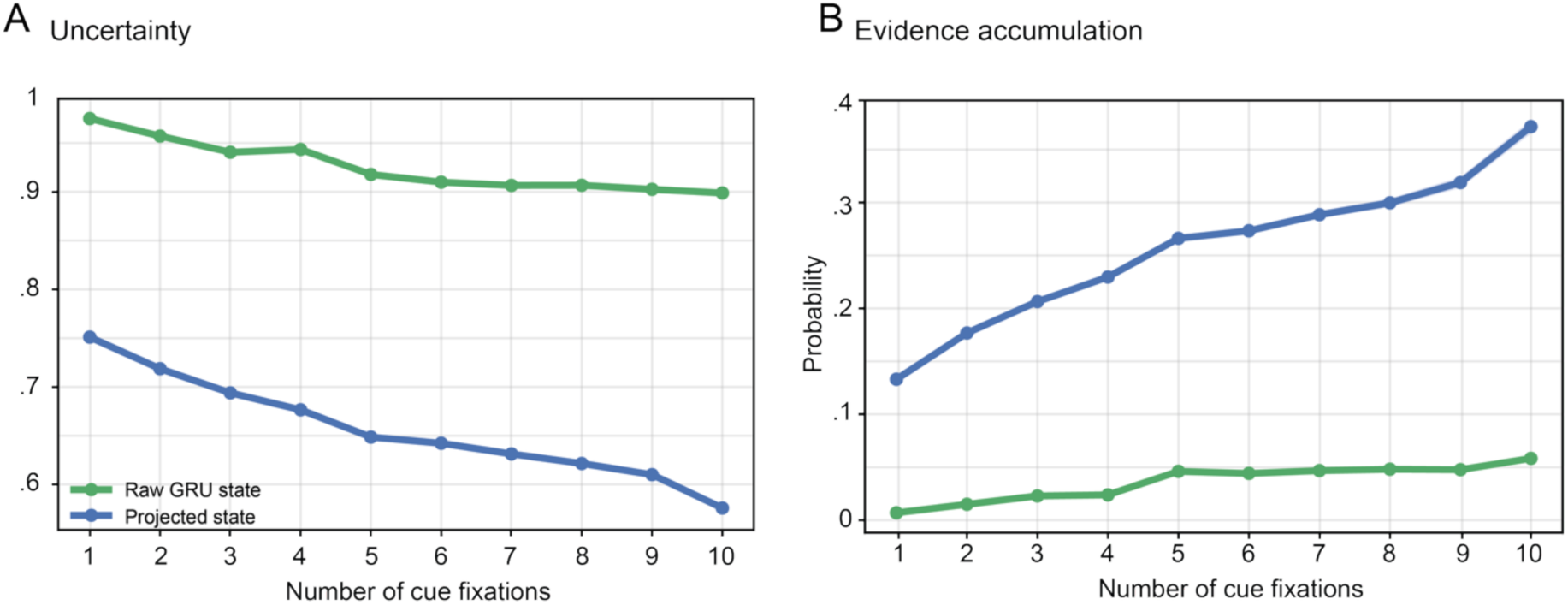
Uncertainty. **A** As additional fixations are accumulated, representational uncertainty progressively decreases both in raw GRU state (green) and in the projected state (blue). B Evidence accumulation also increased across fixations in both raw GRU state (green) and in projected state (blue), however, more steep in the projected state.

**Supplemental Figure 2.**
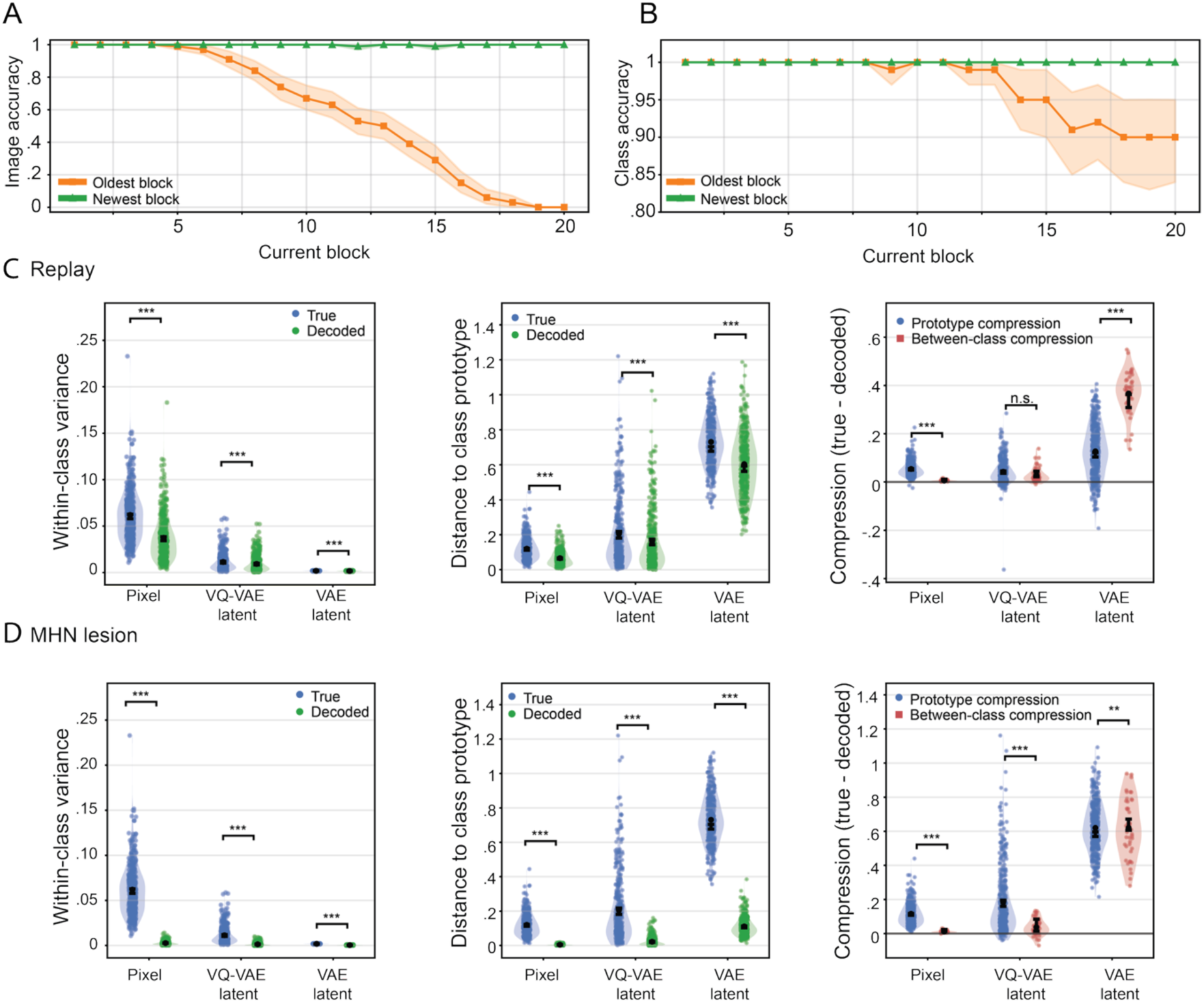
Degradation of compressed memory representations preferentially preserves semantic over item-specific information. **A** Progressive weakening of associative-memory strengths produced a pronounced temporal gradient in exact-image retrieval. Accuracy declined most strongly for the oldest memories, whereas recently encoded memories remained highly retrievable. Lines show observed accuracy and shaded regions show class-stratified bootstrap 95% confidence intervals. B Semantic category retrieval remained comparatively robust during memory decay, including for the oldest memories. Lines show observed accuracy and shaded regions show class-stratified bootstrap 95% confidence intervals. C Replay reorganised representational geometry in a space-dependent manner. Replayed images were reconstructed by a cortical VAE and evaluated in pixel space, an independent frozen VQ-VAE space, and the replay-trained VAE latent space. Replay reduced withín-class variance and prototype distance in all spaces. Prototype compression exceeded between-class compression in pixel space; the same numerical tendency was smaller in frozen VQ-VAE space, whereas the replay-trained VAE latent space showed broader between-class contraction. D When MHN-mediated retrieval was bypassed, cortical-only reconstructions showed pronounced reductions in within-class variance and movement towards class prototypes. Prototype compression exceeded between-class compression in pixel and frozen VQ-VAE spaces. In the replay-trained VAE latent space, both prototype and between-class compression were substantial. In C-D, violins and points show image-level or class-centroid-pair distributions; black markers show observed estimates and vertical bars show bootstrap 95% confidence intervals. Significance brackets indicate bootstrap contrasts: *p<.05, ** p <.01,*** p <.001; n.s„ not significant.

**Supplemental Figure 3.**
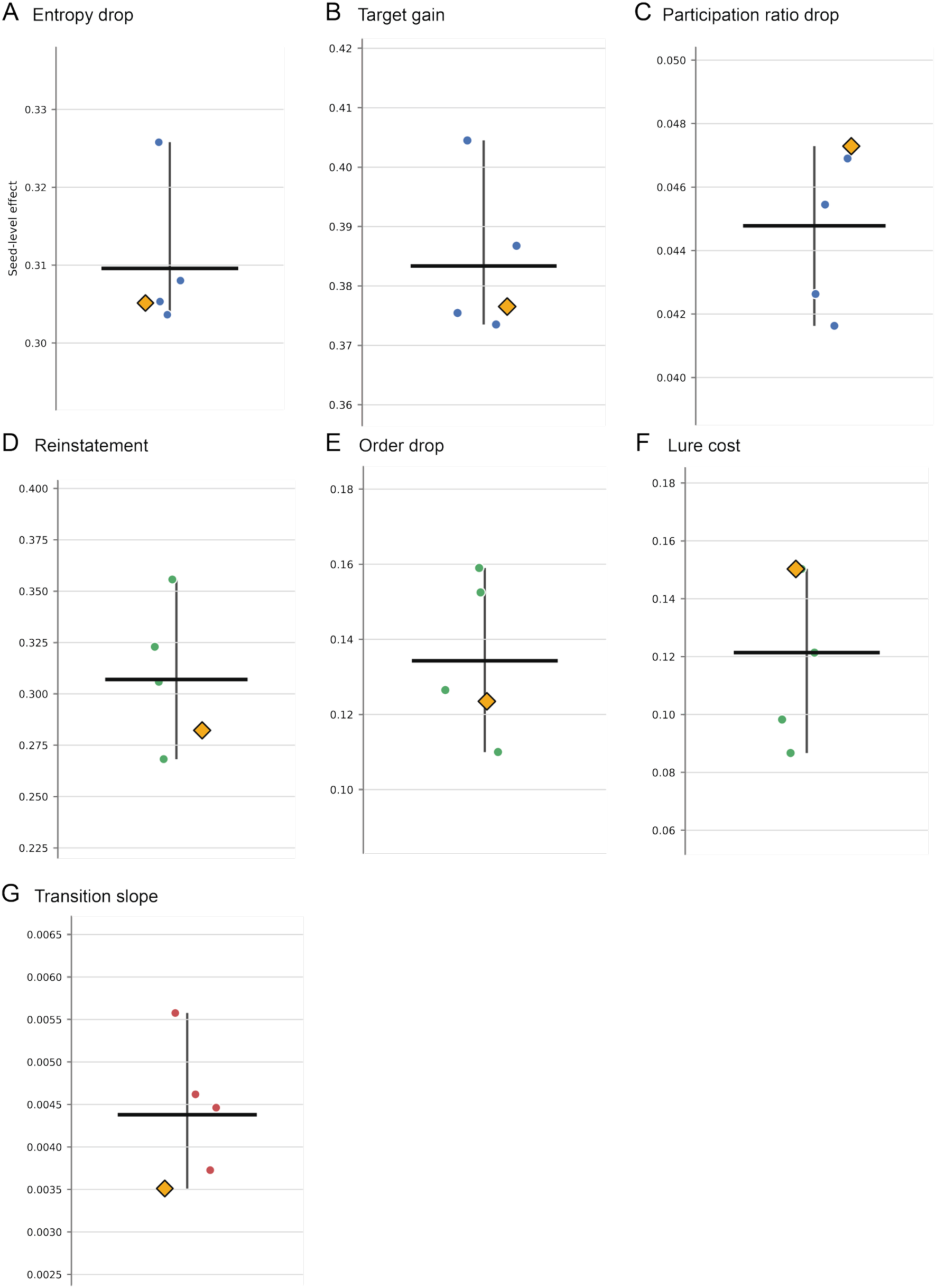
Core effects across independently trained seeds. **A** Entropy reduction. B Target-evidence gain. C Participation-ratio reduction. D Reinstatement slope. E Reversed-order cost. F High-overlap lure cost. G Relational-transition slope. Points denote seeds; gold diamonds mark the paper seed, black bars the mean, and lines the range.

